# Structural modelling uncovers diverse predicted transcriptional and post-transcriptional modulators among type III effectors of symbiotic Rhizobia

**DOI:** 10.64898/2026.07.12.738046

**Authors:** Albin Teulet, Sebastian Schornack

## Abstract

Rhizobia are soil bacteria that establish nitrogen-fixing symbioses with legumes. While many rhizobia use a Type III Secretion System to deliver “Nodulation Outer Protein” (Nop) effectors, some uniquely use these proteins to initiate nodule organogenesis, bypassing classical signalling. The molecular functions of these effectors remain largely unknown due to extreme sequence divergence. Using AlphaFold2-mediated structural proteomics, we identified a modular architecture in rhizobial effectors composed of 22 distinct structural units. We reveal that many Nop effectors are cryptic transcriptional or post-transcriptional regulators, harbouring unrecognised nucleic acid–binding modules and RNA-dependent RNA polymerase domains. Crucially, these modules are conserved in specific plant pathogens, such as gall-inducing *Pantoea*, where our predicted structural units align with experimentally validated DNA-binding domains. Furthermore, we discovered the BPN (B3 and PUA-like nucleic acid binding) domain as a structural mimic of plant B3-domain transcription factors, pointing to a direct mechanism for hijacking legume development. Our findings strongly suggest that rhizobia employ a modular domain-fusion strategy to act as direct genetic modulators, uncovering a conserved mechanism used by both symbionts and pathogens to hijack host developmental programmes.

## Introduction

Effectors are a hallmark of host-pathogen interactions. They are encoded in the microbe and are translocated into the host tissues where they interfere with host processes. Studying effectors has advanced our understanding of pathogen strategies^1^, supported the generation of disease resistant plants^2^ and has fuelled bioengineering approaches^3^. In Gram-negative pathogenic bacteria, the Type III secretion system (T3SS) is a crucial pathogenicity determinant. It forms a multiprotein complex in the bacterial double membrane and architecturally analogous to a syringe, facilitating the direct translocation of type III effector proteins (T3Es) into host cells^4^. Once inside, effectors target key host processes including protein degradation, trafficking and cytoskeleton dynamics, and gene expression^5^. Indeed, some *Xanthomonas* plant pathogens have evolved an even more sophisticated strategy by employing effectors that directly bind to host DNA to modulate transcriptional programmes. These transcriptional activator-like (TAL) effectors enter the nucleus with the help of nuclear localisation signals (NLS) where they bind to promoters of host susceptibility genes and activate their expression^3,6^.

The Type III Secretion System (T3SS) is utilised not only by pathogenic bacteria but also by soil bacteria within the Rhizobacteria group. These bacteria establish beneficial nitrogen-fixing symbioses within nodules, specialised organs formed by the roots of legume plants^7^. Despite being recognised as symbionts, a T3SS machinery, as well as T3Es, have been identified in several rhizobia genera, including *Rhizobium*, *Sinorhizobium*, *Mesorhizobium*, and, most abundantly, in *Bradyrhizobium*^8,9^. Approximately 20 proteins, called Nodulation Outer Proteins (Nops), are now defined as bona fide effectors, a designation supported by experimental evidence of their secretion through the T3SS machinery^10,11^. The secretion of rhizobia effectors during the symbiotic interaction can have dual outcomes^12^. On one hand, they can facilitate the establishment and maintenance of the symbiosis by suppressing the host plant’s immune responses^13,14^. On the other, they may be detected by the plant’s resistance (R) proteins, thereby eliciting an immune response that impedes the infection process^15,16^.

The knowledge regarding the function of Nops and their roles in symbiotic interactions is limited. However, recent years have seen a surge in studies elucidating their functions. Numerous functional domains have been identified and characterised, facilitating the classification of Nops into functional families. Notably, the NopM family carries an N-terminal Leucine-rich repeat (LRR) and a C-terminal Novel E3 ubiquitin ligase (NEL) domain^17^, the NopD family includes a C-terminal Ubiquitine-like protease (ULP) domain^18,19^, and the NopT family features the C58 cysteine protease domain^20^. Despite these advances, many Rhizobia effectors remain functionally uncharacterised, with primary sequence analysis alone proving insufficient to predict their molecular functions^10,11^.

Recently, several effectors from bacteria of the genus *Bradyrhizobium* have been identified as sufficient to initiate organogenesis processes, resulting in the formation of nodule-like structures on *Aeschynomene* and soybean legume plants. Among these effectors are ErnA from *Bradyrhizobium vignae* ORS3257^21^, Bel2-5 from *B. elkanii* USDA61^19^, and Sup3 from *Bradyrhizobium* sp. NAS96.2^22^. These ET-Nod (Effectors Triggering Nodulation) effectors^23^, exhibit significant sequence variability, yet all harbour NLS motifs and localise to plant cell nuclei once transiently expressed in tobacco cells^19,21,22^. Moreover, ErnA associates with nucleic acids *in planta*^21^, suggesting that it may directly interact with DNA to modulate plant gene expression and developmental programmes. Yet, DNA or RNA-binding domains have not been reported for effectors of bacterial symbionts.

In this computational study, we aimed to gain deeper insight into the molecular functions of rhizobial type III effectors. We combined several complementary bioinformatic approaches integrating sequence and structural model similarity analyses. Our investigations revealed that many effectors display a modular architecture composed of multiple structurally discrete domains, several of which show similarities to known enzymatic domains. Surprisingly, we also identified three distinct potential nucleic acid–binding domains previously unrecognised in rhizobacterial effectors often co-occurring with predicted transcriptional activation domains. In certain effectors, the presence of potential sequence-specific RNA-interaction domains is consistently associated with predicted RNA-dependent RNA polymerase modules, which are implicated in the biogenesis of small double-stranded RNAs.

Our findings uncover a previously unrecognised group of rhizobial effectors that may act as transcriptional or post-transcriptional modulators in plants, and strongly suggest that nucleic acid interactions represent a common strategy used by symbiotic bacteria to manipulate plant host cellular programmes.

## Results

### Shared structural units underpin diverse Nop effector architectures

We assembled a dataset of 33 Nodulation Outer Protein (Nop) effectors from *Bradyrhizobium*, *Sinorhizobium*, and *Mesorhizobium* spp., representing the currently described diversity of rhizobial type III-secreted effectors (S1 Fig., S1 Table). Sequence-based domain prediction using InterProScan identified 13 distinct domains in 18 effectors, including enzymatic domains such as acetyltransferases, C48 and C55 peptidases, C58 cysteine protease, shikimate kinases and NEL-type E3 ubiquitin ligase, as well as integrase and leucine-rich repeat domains (S1 Fig., S2 Table). In several cases (e.g., NopT^NGR234^, NopM^NGR234^, NopF^USDA61^, NopJ^NGR234^, NopAC^ORS3257^, NopAD^ORS3257^, GunA^USDA257^), these domains spanned most of the protein length, whereas others, such as Ubiquitine-like protease (ULP) and shikimate kinase domains, were restricted to short C-terminal regions of otherwise unannotated proteins (S1 Fig., S2 Table).

No known domains could be assigned to the remaining 15 Nop effectors, underscoring the limitations of sequence-based functional inference. To improve effector annotations, we pursued a structural modelling and comparison approach. We generated structural models for all 33 Nops using AlphaFold2^24^ and selected, for each effector, the highest-confidence model among the five predictions based on the global pLDDT score^25^. Overall model confidence varied widely across the dataset (S2A Fig.), with approximately half of the proteins exhibiting a low global pLDDT score ≤ 70, consistent with the presence of long unstructured regions likely corresponding to intrinsically disordered segments.

To address this, we examined the local pLDDT profiles of each best model and extracted well modelled regions with pLDDT scores ≥ 70. This approach identified 96 structural regions across the 33 Nop structural models^25^ (S2B, S3, S4A Fig., S2 Table). Structural comparison and clustering of these regions using a TM-score cutoff ≥ 0.5 identified 13 structural communities with at least two members and 9 singletons across the Nop effector set (Fig. 1A,S4B Fig., S3 and S4 Tables). In total, 22 distinct structural units (SUs) were defined. These SUs occur singly, in tandem repeats, or in modular combinations within individual Nop proteins. Among them, 10 SUs correspond to domains that could be identified by sequence-based annotation and already have a putative function assigned (S1 Fig., S2 Table).

**Fig. 1.**
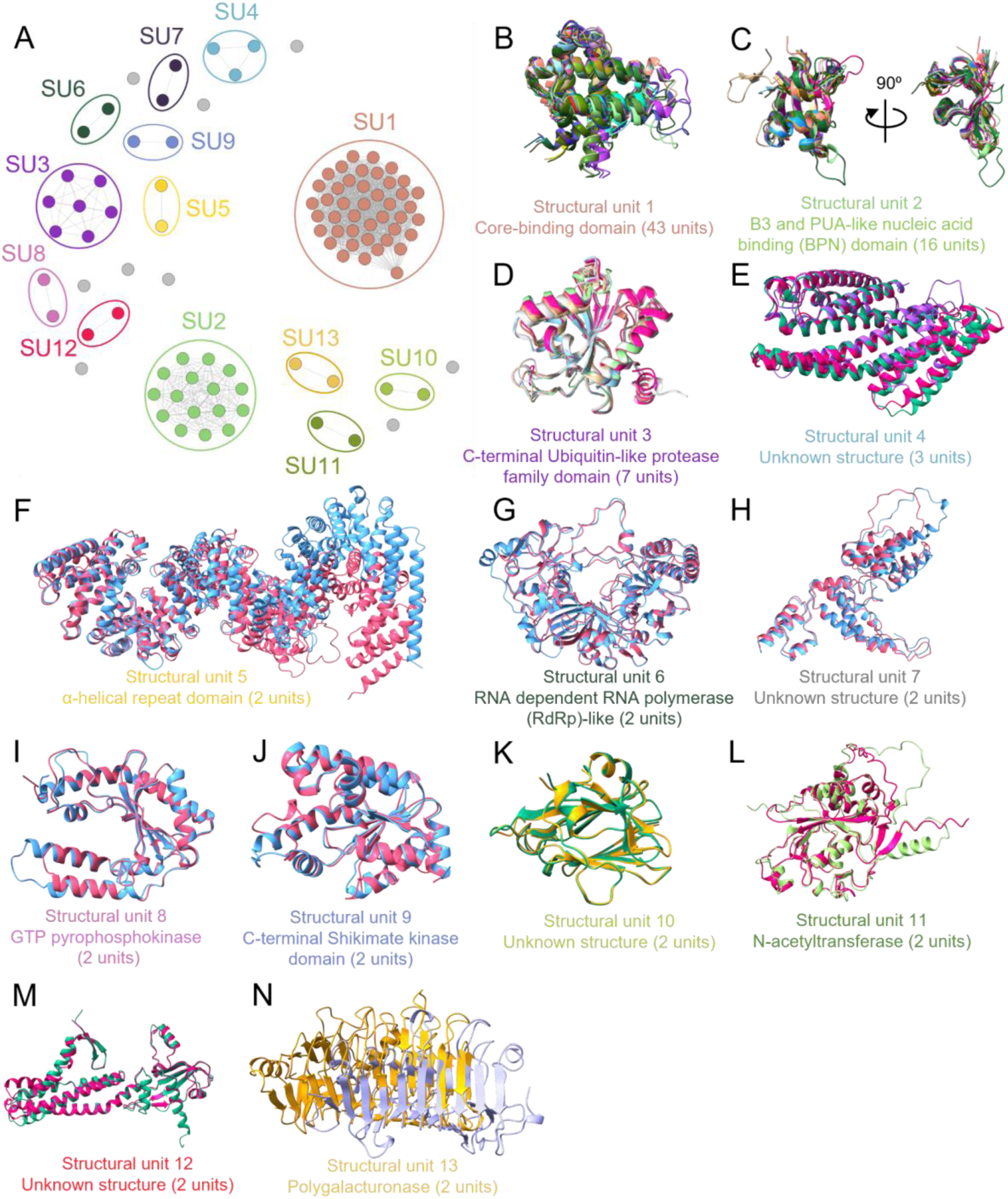
Network-based identification of structural units (SUs) present in multiple in Nop effectors (A) Structural similarity network of the 96 structural units (SU) extracted from AlphaFold models of the 33 Nop effectors. Each node represents one region, and edges connect pairs of regions that are structurally similar with a TM-score above 0.5. Coloured circles delimit the 13 communities containing at least two members (SU1–SU13). The remaining grey nodes correspond to the 9 singleton units. (B–N) Overlays of all structural regions belonging to each community (SU1–SU13). The putative fold or function inferred from Foldseek analysis is indicated below each panel, with the number of occurrences in each community given in parentheses

To validate our structural modelling and segmentation strategy, we assessed whether AlphaFold2 recaptures experimentally determined folds. For the 10 sequence-annotated SUs, structural superpositions against available crystallographic references showed moderate to high similarity, with root mean square deviations (RMSD) < 3 Å and TM scores ≥ 0.5 (S5 Fig.). These results indicate that our pipeline robustly delineates structural units that correspond to experimentally resolved folds in rhizobial effectors.

Finally, to infer possible functions of the remaining 12 SUs without sequence matches, we conducted Foldseek-guided structural similarity searches^26^ against the AlphaFold-predicted proteomes, including databases generated from UniProt50 and Swiss-Prot, as well as the PDB100 structural database. This analysis provided functional annotations for 6 of the 12 remaining SUs without sequence matches (Fig. 1B, 1C, 1F, 1G, 1I, 1L, S6 Fig., S5 Table), notably revealing similarity to enzymatic and ligand-binding domains, while the remaining 6 could not be confidently assigned a putative function.

### Structural modelling reveals new enzymatic domains in Nop effectors

Our structure similarity searches found three matches to enzymes that were not identifiable through their sequence. Among these, the SU6 present in Mlr6331^MAFF303099^ and in Mlr6361^MAFF303099^ is similar to two-barrel polymerases (S5 Table). In particular, SU6 aligns with the fungal RNA-dependent RNA polymerases (RdRPs) QDE-1 from *Thielavia terrestris* (PDB: 5FSZ, TM-score = 0.555) and *Neurospora crassa* (PDB: 7Y7R, TM-score = 0.556), as well as with Rdp1 from *Schizosaccharomyces pombe* (AlphaFold model AF-O14227-F1, TM-score = 0.524) (Fig. 2A, S7A Fig.). SU6 adopts the characteristic two-barrel architecture of this polymerase family, consisting of a U-shaped arrangement with a double φ-β-barrel catalytic core positioned at the base of the unit^27,28^. In *S. pombe*, Rdp1 is a key component of the RNA interference (RNAi) pathway, essential for the biogenesis of small interfering RNAs (siRNAs), and required for both post-transcriptional and transcriptional gene silencing^29^. Structural alignment confirms the conservation of the catalytic domain within SU6, including the critical aspartate residue necessary for RdRP activity^30^ (Fig. 2B, S8 Fig.).

**Fig. 2.**
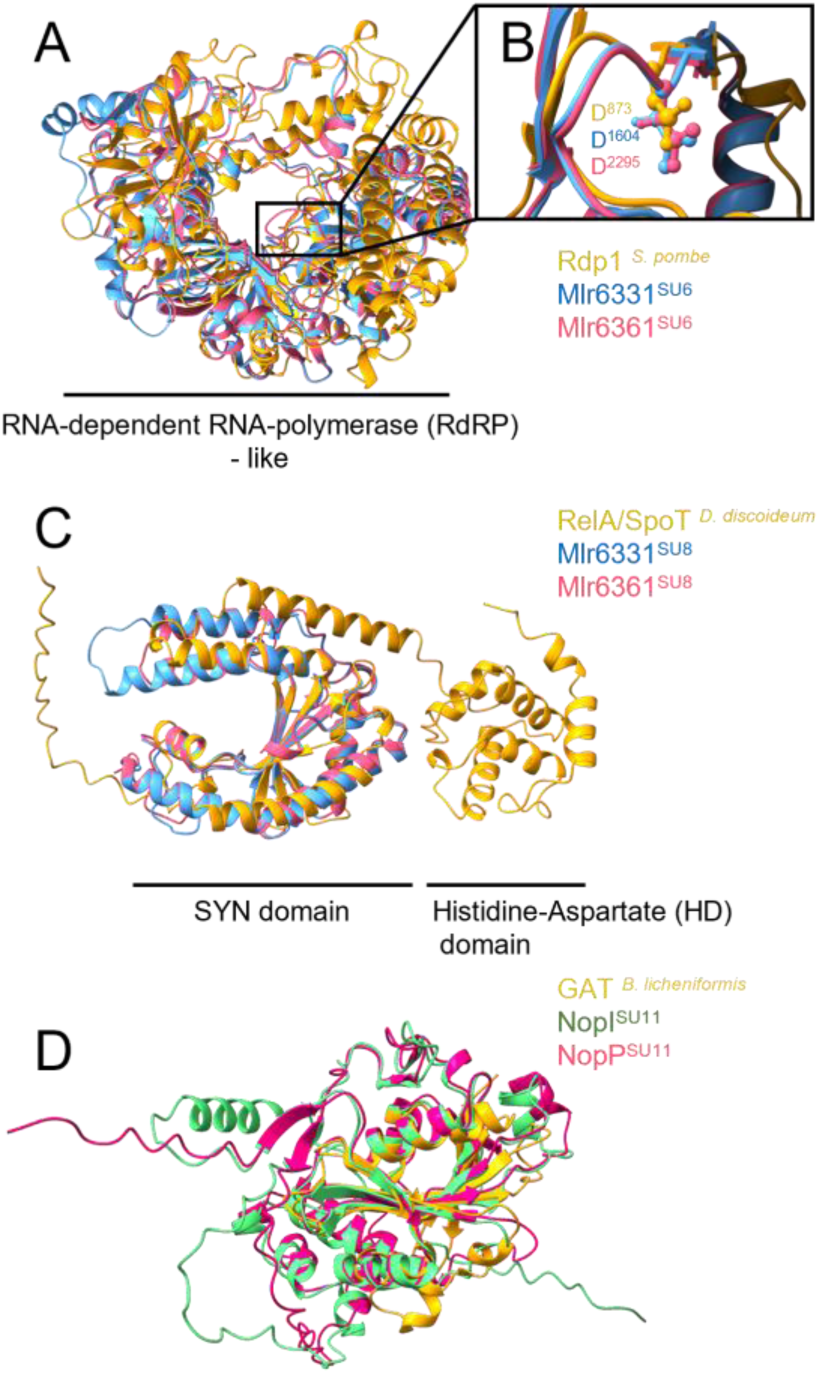
Cryptic enzymatic architectures revealed in Nop effectors Three structural units adopt cryptic enzymatic folds despite the absence of detectable global sequence similarity. (A) SU6, conserved in Mlr6331^MAFF303099^ and Mlr6361^MAFF303099^, shows structural similarity to the RNA-dependent RNA polymerase Rdp1 from *Schizosaccharomyces pombe* (AF-O14227-F1). (B) Local sequence similarity is detectable around the catalytic site of the polymerase, where the key catalytic aspartate residue is conserved. (C) SU8 adopts a fold similar to SYN synthetase domain of GTP pyrophosphokinases, including the RelA/SpoT domain-containing protein from *Dictyostelium discoideum* (AF-Q55FF4-F1). (D) The two effectors NopP^USDA110^ and NopI^HH103^ share SU11, which displays structural similarity to glyphosate N-acetyltransferase (GAT) from *Bacillus licheniformis* (PDB: 2JDD).

Upstream of this SU6 in Mlr6331^MAFF303099^ and in Mlr6361^MAFF303099^, SU8 displays structural similarity to the synthetase (SYN) domain of several GTP pyrophosphokinases, including ppGpp synthetase (RelA) and ppGpp synthetase/hydrolase (SpoT) (Fig. 2C, S5 Table). In prokaryotes, ppGpp serves as a second messenger involved in multiple cellular processes, including the regulation of transcription elongation through its interaction with RNA polymerase^31^.

Finally, Foldseek-guided structural comparison of SU11 supports the assignment of putative enzymatic functions to NopP from *B. diazoefficiens* USDA110 and NopI from *M. loti* HH103, both of which harbour this structural unit that exhibits high-confidence similarity to N-acetyltransferase family proteins across multiple taxa (Fig. 2D, S5 Table).

### Cryptic structural modules with predicted nucleic acid–binding functions are widespread in Nop effectors

Besides new domains resembling enzymes, we uncovered previously unrecognised regions within Nop effectors that display strong similarity to known nucleic acid binding domains. Among the identified structural units, SU1 is the most abundant and present in 13 of the 33 effectors analysed. SU1 shows strong structural similarity to the core-binding (CB) domain (IPR044068) of tyrosine-type site-specific recombinases (Fig. 3A and 3B, S5 Table). In recombinases, the CB domain participates in essential DNA-associated processes, including supercoil relaxation, plasmid and chromosomal partitioning, and phage integration/excision^32^. CB domains interact primarily with the major groove of DNA and contribute to both DNA recognition and protein–protein interactions^33^. Although the SU1 units identified in Nop effectors are structurally most similar to bacterial tyrosine recombinases, they do not co-occur with recombinase catalytic domains. Instead, SU1 is present in one to six copies per effector and frequently co-occurs with SU2 (BPN domain, see below) and/or SU3 (Ubiquitine-like protease) domains in a modular architecture (S1 Fig.). Evolutionary analysis of the 43 SU1 sequences revealed extensive amino acid divergence (S9A Fig.), yet conservation mapping on the SU1 structure identified a restricted set of conserved hydrophobic residues that cluster into an internal structural core and form the most evolutionarily constrained regions of the fold (S9B Fig.). This alignment between structural stability, high pLDDT scores, and evolutionary constraint provides strong support for the functional relevance of these previously unrecognised domains.

**Fig. 3.**
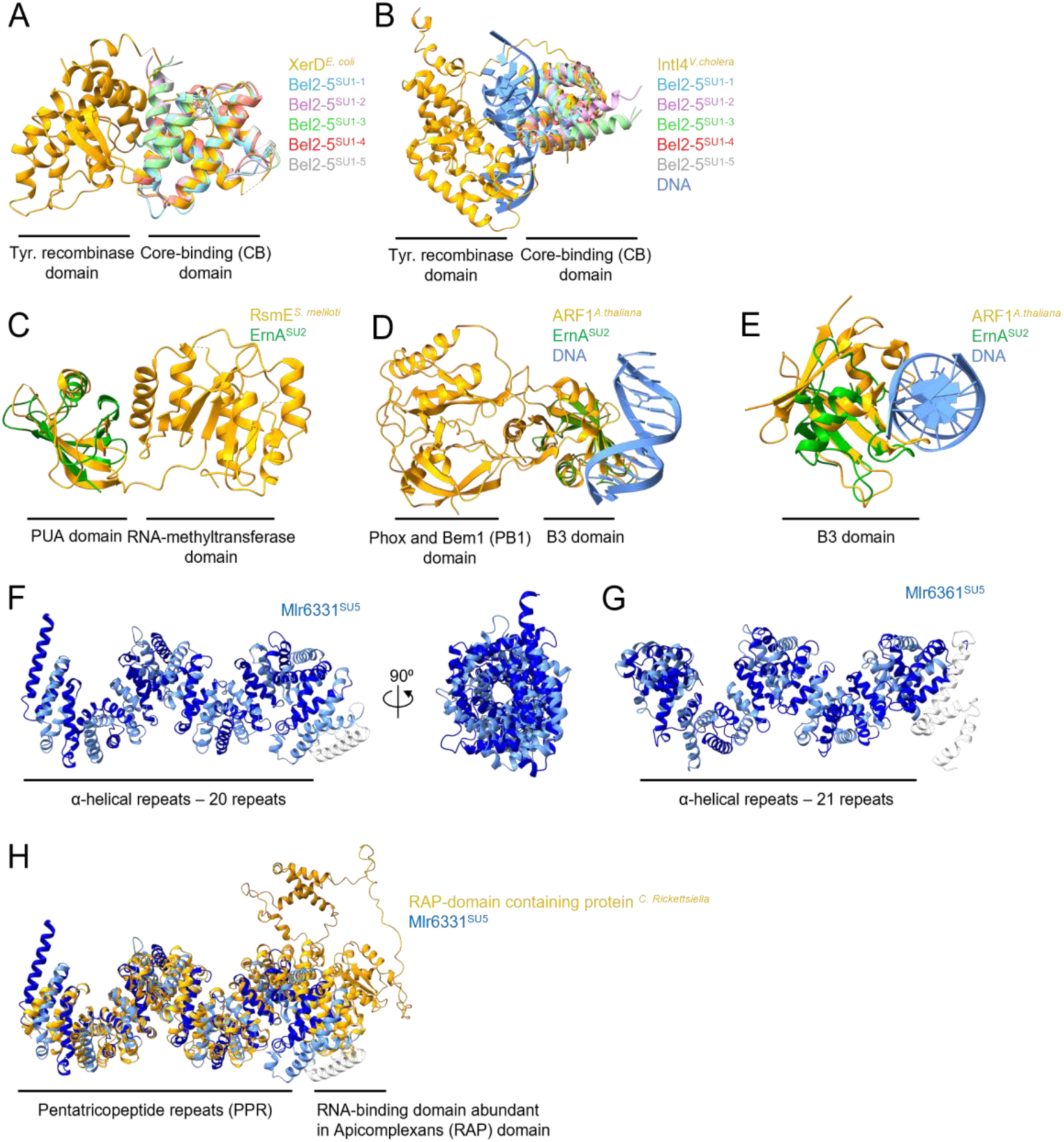
Putative nucleic acid–binding folds in Nop structural units. Structural comparisons of Nop structural units SU1, SU2 and SU5 with known nucleic acid–binding proteins. Effector structural units are coloured and reference proteins are shown in yellow. (A,B) The five SU1 copies from Bel2-5^USDA61^ are superimposed on the core DNA-binding domains of the tyrosine recombinases XerD from *Escherichia coli* (PDB 1A0P; A) and IntI4 from *Vibrio cholerae* (PDB 2A3V; B). (C–E) The BPN (SU2) domain of ErnA^OSR3257^ structurally aligns with the pseudouridine synthase and archaeosine transglycosylase (PUA) RNA-binding domain of RsmE from *Sinorhizobium meliloti* (PDB 4J3C; C) and with the B3 DNA-binding domains of ARF1 (PDB 4LDX; D) and ARF19 (AlphaFold accession: AF-Q8RYC8-F1; E) from *Arabidopsis thaliana*. (F,G) SU5 from Mlr6331^MAFF303099^ (F) and Mlr6361^MAFF303099^ (G) adopts a multi-helical α-solenoid architecture composed of tandem repeats. (H) SU5 from Mlr6331^MAFF303099^ overlaid with the PPR domain of a RAP-domain–containing protein from *Candidatus Rickettsiella isopodorum* (AlphaFold accession: AF-A0A1J8P2S6-F1).

The SU5 structural unit is found exclusively in Mlr6331^MAFF303099^ and Mlr6361^MAFF303099^. SU5 adopts a multi-helical α-solenoid architecture composed of tandem repeats arranged in a right-handed superhelix. Each repeat spans about 40-45 amino acids, with approximately 20 repeats in Mlr6331^MAFF303099^ and 21 in Mlr6361^MAFF303099^ (Fig. 1F, Fig 3. F and 3G). Foldseek-based searches identified numerous structurally similar proteins, most of which are annotated as pentatricopeptide repeat (PPR)-containing proteins in plants, indicating that SU5 is closely related to PPR-like α-helical repeat folds (S5 Table). Pentatricopeptide repeat proteins are sequence-specific RNA-binding proteins in which individual repeats contribute to the recognition of defined nucleotide positions along the RNA^34^. In Mlr6331^MAFF303099^ and Mlr6361^MAFF303099^, SU5 shows the highest structural similarity to the α-helical repeat region of a RNA-binding domain abundant in apicomplexans (RAP) domain-containing protein from *Candidatus Rickettsiella isopodorum*, with TM-scores of 0.687 and 0.703, respectively. Yet, SU5 is not linked to a RAP domain in these effectors but upstream of the RdRP-like SU6 (S1 Fig., S5 Table). As RAP proteins and related PPR repeat proteins have been implicated in RNA-binding, It is tempting to speculate that SU5 and RdRP-like SU6 operate as a functional module in which RNA recognition by SU5 could guide SU6-mediated RNA synthesis.

### SU2 defines a novel structural domain with nucleic acid–binding features in Nop effectors

SU2 represents the second most abundant structural unit in our dataset, with 16 occurrences distributed across 10 Nop effectors. This unit is consistently located in the C-terminal region and exhibits a conserved modular organisation with other structural units (S1 Fig.). SU2 frequently co-occurs with SU1 (Core-DNA binding domain), which is always N-terminal to SU2, and when present alongside SU3 (ULP-like domain), SU2 is positioned between SU1 and SU3. The number of SU2 copies per effector varies, occurring as a single unit (e.g., NopL^NGR234^), as tandem duplications (e.g., NopAB^ORS3257^, InnB^USDA61^), or as triplet arrays (e.g., NopAR^ORS3257^). Notably, five of the six effectors previously characterised as ET-Nods also harbour SU2, indicating that this unit is recurrent among this effector family (S1 Fig., S2 Table). In ErnA^ORS3257^, the C-terminal SU2 harbours the sequence motifs essential for its activity^22^ (S10 Fig.).

Again, SU2 is structurally, but not sequence-conserved (S11A Fig.). Mapping of evolutionary conservation scores onto SU2 models showed elevated conservation in a structural core supporting that SU2 represents a genuine and functionally constrained structural unit (S11B Fig.).

Foldseek searches for SU2 in the AlphaFold databases returned multiple and varied hits across SU2 occurrences and TM-scores were all below the 0.5 confidence threshold, preventing a robust assignment of a putative function (S4 Table). To complement this analysis, we then performed a second search for SU2-similar occurrences using the DALI server^35^ against the PDB database and the full set of AlphaFold-predicted *Arabidopsis* proteome models (S6 Table). This survey showed that most SU2 hits exhibit structural similarity to the bacterial RNA methyltransferase RsmE (PDB: 4J3C), specifically within the pseudouridine synthase and archaeosine transglycosylase (PUA) RNA-binding domain (IPR002478, TM- score = 0.563) (Fig. 3C, S6B Fig.). SU2 also displays structural resemblance to B3 domains (IPR003340), a plant-specific DNA-binding fold found in multiple plant transcription factor families, including Auxin Response Factors (ARFs)^36,37^, Reproductive Meristem (REMs)^38^, Related to ABI3/VP1 (RAV)^39^, and LEC2/ABI3/VAL proteins^40^ with *Arabidopsis* ARF19 (TM-score = 0.509) as the best match, particularly across the β strands known to mediate DNA sequence-specific recognition and positioned laterally to the major groove of the DNA target^41^ (Fig. 3D and 3E, S7B Fig.). The high degree of fold preservation between SU2 and the plant B3 domain suggests a case of molecular mimicry, potentially allowing rhizobial effectors to compete for or occupy the same DNA-binding sites as host Auxin Response Factors (ARFs).

Since SU2 exhibits structural features of both PUA and B3 domains yet cannot be reliably assigned to either family, we refer to it as a B3 and PUA-like nucleic acid binding (BPN) domain.

Taken together, these data reveal that many rhizobial type III effectors harbour structurally distinct units similar to nucleic acid–binding domains, suggesting that the ability to interact with DNA and/or RNA is a recurrent and potentially central feature of Nop effector families.

### Nop effectors commonly harbour predicted nuclear localisation signals

The widespread occurrence of putative nucleic acid–binding structural units among Nop effectors prompted us to examine whether these effectors are able to localize to the host nucleus. Nuclear localisation has previously been reported for 5 of the 33 rhizobial type III effectors examined in this study, including the ET-Nods ErnA^ORS3257^ ^21^, Sup3^NAS96.2^ ^22^ and Bel2-5^USDA61^ ^19^, as well as NopD^XS1150^ ^18^ and NopL^NG234^ ^13^, suggesting that nuclear targeting may be a recurrent feature of rhizobial effector families. LOCALIZER^42^ predicted between one to four NLS motifs in 16 of the 33 Nop effectors (Fig. 4A, S1 Fig., S7 Table). Notably, 14 of the 16 effectors with a predicted NLS also contain at least one of the three structural units with nucleic acid binding similarity, supporting a strong association between nuclear targeting and nucleic acid–binding potential in Nop effectors. Interestingly, despite being reported to translocate into the plant cell nucleus, no NLS was detected in the effectors NopL^NGR234^ and NopD^XS1150^.

**Fig. 4.**
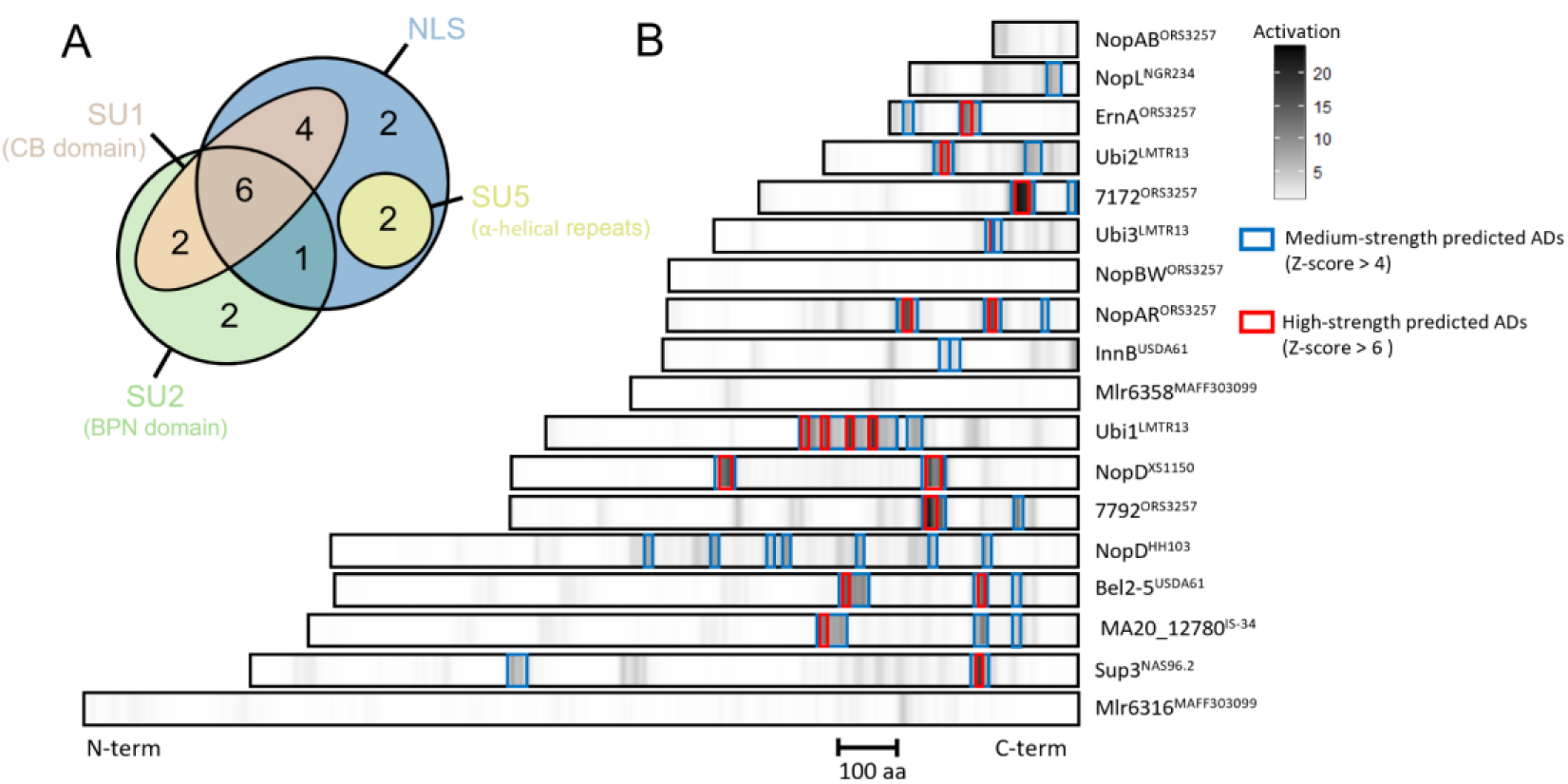
Candidate transcriptional activation domains are present in most of the studied rhizobial effectors with predicted nucleic-acid binding units. (A) Venn diagram showing the overlap between Nop effectors carrying putative nucleic acid–binding structural units SU1 (CB domain), SU2 (BPN domain) and SU5 (α-helical repeats) and those with a nuclear localisation signal (NLS) predicted by LOCALIZER. Most proteins containing SU1, SU2 or SU5 also harbour a predicted NLS. (B) Heatmap of PADDLE-predicted transcriptional activation scores along Nop effector sequences, plotted from N- to C-terminus and ordered by protein length (scale bar, 100 aa). Darker shades indicate higher predicted activation. Blue boxes highlight medium-strength activation domains (Z-score ≥ 4), and red boxes indicate high-strength activation domains (Z-score ≥ 6).

### Conservation of predicted transcriptional activation domains in Nop effector families

To assess whether Nop effectors could be transcription factors, we searched for the presence of transcriptional activation domains (ADs) using PADDLE (Predictor of Activation Domains using Deep Learning)^43^. The analysis was performed on all effectors containing the SU1 (core-binding) or SU2 (BPN) domains, as well as on NopBW^ORS3257^ and Mlr6358^MAFF303099^, which carry an NLS but lack any predicted nucleic acid–binding domain.

Among the 18 effectors analysed, PADDLE predicted at least one transcriptional AD in 14 of the 18 effectors analysed, with 11 showing high-confidence prediction scores (Z-score ≥ 6) (Fig. 3B, S8 Table). No AD was detected in the two control effectors NopBW^ORS3257^ and Mlr6358^MAFF303099^, both of which lack predicted nucleic acid–binding domains. To complement this analysis, we also applied TADA (Transcriptional Activation Domain Activity)^44^, a deep learning predictor trained specifically on plant activation domains. All sites predicted by PADDLE were likewise identified by TADA (Fig. S12, S8 Table), yielding a common set of 27 robust candidate activation domains across the 14 effectors with positive predictions (S1 Fig.).

To evaluate whether these ADs are restricted to individual variants or represent conserved features within each effector family, we next identified homologues of all analysed effectors across a dataset of 654 rhizobial proteomes spanning the *Bradyrhizobium*, *Sinorhizobium*, and *Mesorhizobium* genera. This search recovered a total of 643 homologous proteins. Detection of ADs with PADDLE across these homologues showed that medium-strength activation domains (Z-score ≥ 4) were predicted in over 75% of the sequences analysed (S13 Fig.). Predicted ADs were less frequent in InnB homologues (65%) and Sup3 homologues (45%). No activation domains were predicted in NopAB^ORS3257^ or Mlr6316^MAFF303099^, but PADDLE detected ADs in 45% and 35% of their homologues, respectively.

Thus, predicted activation domains are broadly found across most homologues in almost all of effector families analysed and many Nop effector families combine key features characteristic of transcriptional regulators (NLSs, predicted nucleic acid–binding domains, and ADs) required to potentially modulate plant gene expression.

### Limited conservation of Nop structural units across bacterial type III effectors

To assess how widely the 22 SUs identified in rhizobial effectors are conserved across bacterial type III effectors, we assembled a dataset of 1,210 type III effector variants reported in the literature (S9 Table). This dataset included 921 effector variants from 11 genera of plant pathogenic bacteria and 287 variants from 13 genera of bacteria that infect animals. Structural models were predicted for all variants, and each model was screened for the presence of the 22 rhizobial SUs by structural similarity analysis^25^ (S10 Table). Among the 1,210 effectors examined, only 93 hits matching 73 effectors had any of the SUs described here (Fig. 5A, S11 Table). The most frequently detected units were SU20 (acetyltransferase; 27 hits) and SU21 (C58 peptidase; 22 hits), which include members of the two widespread type III effector families, YopJ/HopZ and YopT/AvrPphB, respectively. In contrast, eleven SUs showed no detectable matches outside Rhizobia, including SU2 (BPN domain), the second most abundant unit in Nop effectors (Fig. 5A, S10 Table).

**Fig. 5.**
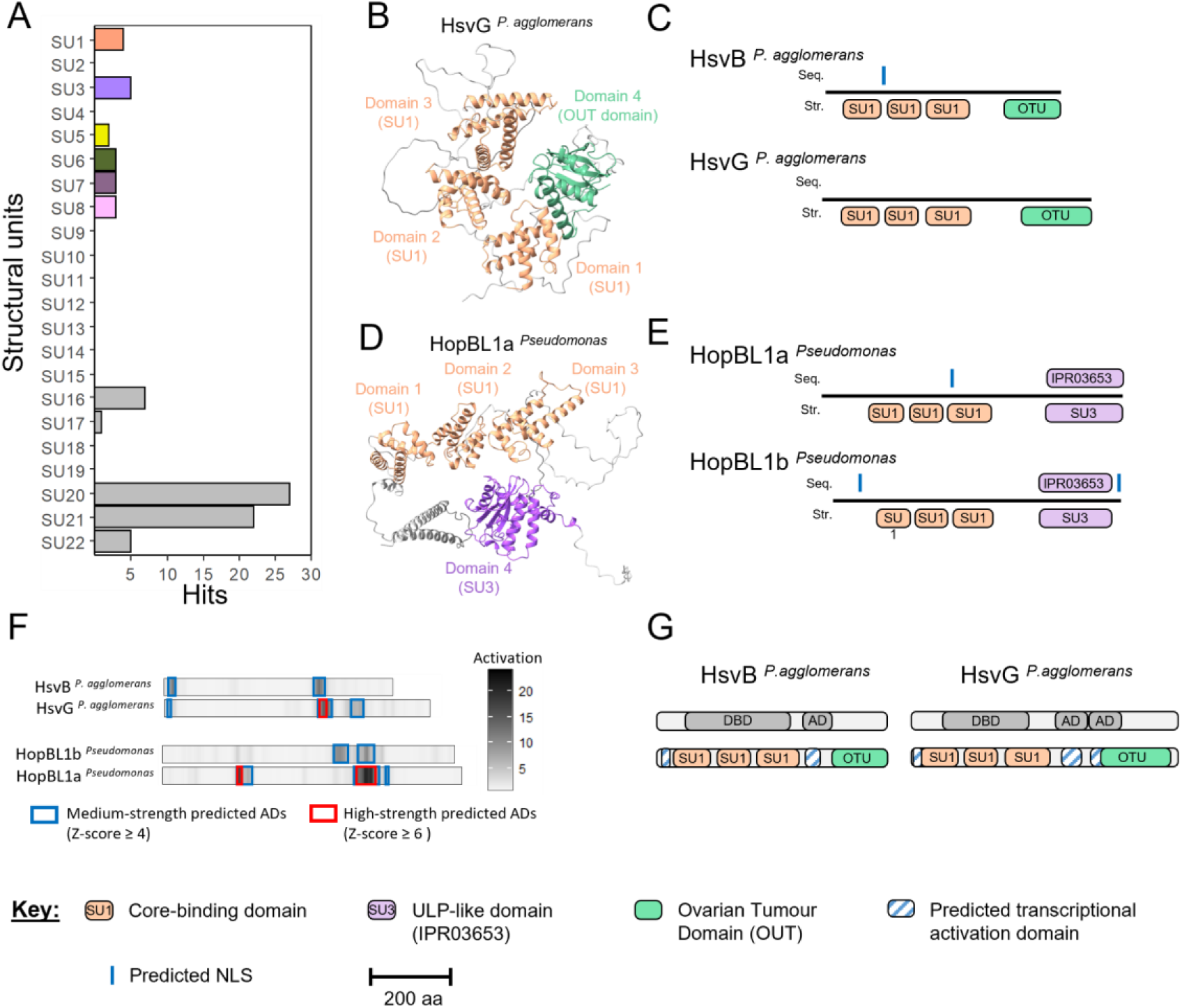
Distribution of rhizobial structural units in non-rhizobial T3Es (A) Number of hits for each structural unit detected by Foldseek in a dataset of 1,210 AlphaFold models of type III effector variants from plant-pathogenic and animal-associated bacteria. Most structural units defined in rhizobial Nops are absent or detected only rarely in this dataset. (B,D) AlphaFold models of HsvG*^P. agglomerans^* and HopBL1a*^Pseudomonas^*, respectively, segmented into four structural units based on pLDDT. In both proteins, the three N-terminal units are structurally identical to SU1, whereas the C-terminal unit corresponds to an OTU-like domain in HsvG*^P. agglomerans^* and to SU3, an ULP-like domain, in HopBL1a*^Pseudomonas^*. (C,E) Linear domain organisation of HsvG*^P. agglomerans^*/HsvB*^P. agglomerans^* and HopBL1a*^Pseudomonas^*/HopBL1b*^Pseudomonas^*, respectively, with sequence-based annotations (seq.) shown above each protein bar and structure-based annotations (str.) shown below. (F) Heatmap of PADDLE-predicted transcriptional activation scores along HsvG*^P. agglomerans^*, HsvB*^P. agglomerans^*, HopBL1a*^Pseudomonas^* and HopBL1b*^Pseudomonas^*, plotted from N- to C-terminus and ordered by protein length (scale bar, 100 aa). Darker shades correspond to higher predicted activation. Blue boxes highlight medium-strength activation domains (Z-score ≥ 4), and red boxes indicate high-strength activation domains (Z-score ≥ 6). (G) Comparison of experimentally defined functional domains in HsvG*^P. agglomerans^* and HsvB*^P. agglomerans^* (top) with the domains identified in this study (bottom) shows that the DNA-binding domain (DBD) maps to the N-terminal array of SU1 core-binding domains, whereas the experimentally validated activation domains (ADs) align with regions enriched in predicted activation domains.

The structural units SU5 (α-helical repeats), SU6 (RdRP-like), SU7 and SU8 (GTP pyrophosphokinase) co-occur in three plant pathogen effectors of unknown function, RipS1 and RipS2 from *Ralstonia* and XopAD from *Xanthomonas citri* (S14 Fig.). These proteins adopt a modular arrangement similar to that of the rhizobial effectors Mlr6331^MAFF303099^ and Mlr6361^MAFF303099^, despite low primary sequence identity, which indicates that the resemblance is largely restricted to the overall domain architecture (S13 Fig.). The architectures diverge at the C-terminal region, which corresponds to an SU9 shikimate kinase domain in Mlr6331^MAFF303099^ and Mlr6361^MAFF303099^, but to a second SU8 unit in RipS1, RipS2 and XopAD. The three plant pathogen effectors belong to the SKPW effector family, defined by conserved tandem repeats of approximately 42 amino acids that systematically start with serine, lysine, proline and tryptophan^45^. Such SKPW motifs are also present in Mlr6331^MAFF303099^ and Mlr6361^MAFF303099^, and they overlap with the α-helical repeats that constitute the structural unit SU5 (S13 Fig.). Thus, symbiotic Mesorhizobia encode SKPW effectors formerly only reported from plant pathogens.

Finally, SU1 (Core-binding domain), the most widely structural unit conserved within our rhizobial dataset, was detected in just four plant pathogen effectors. The homologous effectors HsvB and HsvG from *Pantoea agglomerans* and HopBL1a and HopBL1b from *Pseudomonas* all exhibit a modular architecture similar to that of SU1-containing Nop effectors: several nucleic acid-binding domains followed by a domain that can remove post-translational modifications from target proteins (Fig. 5B–5E). HopBL1 variants carry an SU3-like ubiquitin-like protease domain while HsvB and HsvG have a C-terminal Ovarian Tumour (OTU) domain, a fold conserved in several deubiquitinases^46^ (S5 Table). As observed in Nop effectors containing SU1, NLSs were predicted in HsvB, HopBL1a and HopBL1b, and at least one putative activation domain in all four proteins can be predicted by both PADDLE and TADA. (Fig. 5F, S12 Fig., S7 and S8 Table).

HsvB and HsvG are gall-inducing type III effectors that trigger tumour-like outgrowths on *Gypsophila paniculata* following their delivery into plant cells by *Pantoea agglomerans*^47^. Both proteins have been experimentally characterised and shown to enter the plant cell nucleus, bind to DNA and function as transcriptional activators, altering the expression of host genes involved in cell cycle regulation^47,48^. Their experimentally defined DNA-binding and activation domains match our predictions, providing functional support for the structural and functional organisation inferred from our analyses (Fig. 5G).

## Discussion

We systematically examined 33 Nodulation Outer Proteins (Nops) representing the currently described diversity of rhizobial type III effector families, using comprehensive sequence- and structure-based analyses. We uncovered that many Nops display a strikingly modular architecture composed of a limited repertoire of structural units, several of which show similarity to enzymatic domains, or nucleic acid–binding folds co-occurring with predicted transcriptional activation domains. Collectively, these findings suggest that many rhizobial type III effectors may act as transcriptional and post-transcriptional regulators during symbiosis, a function not previously attributed to bacterial symbionts.

The most abundant structural unit, SU1, shows strong similarity to the core DNA-binding (CB) domain of tyrosine-type site-specific recombinases and occurs in up to six tandem copies across otherwise unrelated Nop families (Fig. 1, Fig. 3). The most compelling evidence for a genuine DNA-binding function comes from its conservation in the gall-inducing effectors HsvB and HsvG from *Pantoea agglomerans*, among the best-characterised bacterial transcriptional activators in plant pathology^47,48^. As we demonstrated in our comparative analysis, the predicted SU1 modules map precisely onto the regions of HsvB and HsvG previously defined as essential for promoter binding (Fig. 5G), providing a strong structural and biological anchor for our inference that SU1 operates as a bona fide DNA-binding module in Nop effectors. This structural conservation, despite negligible sequence similarity, underscores the value of structure-guided analysis for revealing shared molecular strategies across phylogenetically distant bacteria.

The second most abundant unit, the BPN domain (SU2), displays structural similarity to both the PUA RNA-binding domain of bacterial RNA methyltransferases and, most strikingly, to the B3 domain, a plant-specific DNA-binding fold found in multiple plant transcription factor families including Auxin Response Factors (ARFs), RAV, REM, and LEC2/ABI3/VAL proteins^36,37,38,39,40^ (Fig. 3D and 3E). The structural overlay between SU2 and the B3 domain is concentrated at the β-strands known to mediate sequence-specific DNA recognition^41^, raising the possibility that BPN-containing effectors compete for or occupy the same DNA-binding sites as host transcription factors. This hypothesis is particularly compelling given that, in *Medicago truncatula*, MtARF2, MtARF3 and MtARF4 act downstream of the common symbiotic signalling pathway (CSSP) to contribute to nodule initiation^49^. Consistent with a direct role for the BPN domain in effector activity, functional mapping of ErnA^ORS3257^ has localised the sequence motifs essential for nodule induction on *Aeschynomene* to the C-terminal SU2 identified here^20,50^ (S10 Fig.).

Beyond transcriptional regulation, the *Mesorhizobium* effectors Mlr6331^MAFF303099^ and Mlr6361^MAFF303099^ reveal a distinct post-transcriptional regulatory logic, combining a predicted PPR-like α-helical repeat array (SU5) immediately upstream of a structural units similar to RdRP two-barrel polymerase domain (SU6) (Fig. 1F and 2A, S1 Fig.). RdRPs are central to small RNA biogenesis and RNA-directed gene silencing in eukaryotes^27^, and the catalytic aspartate required for polymerase activity is structurally conserved in SU6 (Fig. 2B, S8 Fig.). If these modules operate in analogy to their structural homologues, SU5 may recognise specific host RNAs and direct SU6-mediated synthesis of complementary strands, engaging small RNA pathways to influence host gene expression at post-transcriptional level. Notably, an equivalent SU5–SU6 arrangement was identified in the SKWP effectors RipS1, RipS2 and XopAD from the plant pathogens *Ralstonia solanacearum* and *Xanthomonas citri* (S14 Fig., S11 Table), indicating that this domain combination occurs in both symbionts and pathogens. Only two families of type III effectors have so far been demonstrated to directly activate host transcription: TAL effectors from *Xanthomonas*, which combine a central repeat array for base-specific DNA recognition with a C-terminal activation domain^51^, and HsvB/HsvG from *Pantoea*, which bind host promoters and recruit RNA polymerase to induce gall formation^47^. Our analyses indicate that multiple Nop effector families share this same functional organisation, combining predicted nuclear localisation signals, nucleic acid–binding domains and transcriptional activation domains (Fig. 4, S1 Fig., S8 Table), despite being unrelated to TAL or HsvB/HsvG effectors. This convergence on a common regulatory strategy, achieved through distinct structural solutions, supports a model in which direct manipulation of host gene expression represents a repeatedly selected mechanism for bacterial control of plant development.

Within the Nop repertoire, the ET-Nod effectors ErnA^ORS3257^, Sup3^NAS96.2^, Bel2-5^USDA61^ and the Ubi effectors from *Bradyrhizobium icense* LMTR13 trigger nodule or nodule-like organogenesis on legumes^19,21,22^, yet the molecular mechanisms underlying these activities remain largely unresolved. Nodule organogenesis requires extensive transcriptional reprogramming and depends on key regulators of the CSSP^52,53^, and recent work has shown that interaction with plant SUMO peptides is required for ET-Nod–dependent nodule induction across structurally diverse members of this family^22,50,54^. In this study, we show that all characterised ET-Nods harbour predicted domain combinations supporting a scenario in which ET-Nods promote nodulation by directly reprogramming host developmental programmes, possibly by interfacing with SUMO-dependent regulatory circuits and co-opting transcriptional networks normally governed by plant symbiotic regulators such as ARFs.

In summary, our structural analysis reveals that despite extensive sequence diversity, many rhizobial type III effector families are built from a limited set of conserved, combinatorial modules, many of which resemble nucleic acid–binding folds and frequently co-occur with predicted activation domains. These features strongly suggest that Nop effectors have evolved into sophisticated regulators capable of directly manipulating host gene expression during symbiosis. Experimental validation, including structural characterisation of representative modules and functional assays, will now be essential to confirm their molecular activities and biological relevance during symbiosis. Future work should therefore focus on the identification of their direct DNA and RNA targets in planta and for a deeper understanding of how Nop modules cooperate to reshape host regulatory networks. More broadly, our findings point to the emergence of conserved bacterial nuclear regulators as a key step in the evolution of nitrogen-fixing bacteria-plant interactions, and provide a molecular framework for the future engineering of Nop-derived molecules with biotechnological potential.

## Materials and Methods

### Structure prediction and structural similarity network analysis

Large-scale structural model prediction was carried out with ParaFold^55^ to run AlphaFold2^24^ on all effector protein sequences analysed in this study. Five structural models were computed and the top-ranked model (ranked_0, according to the pLDDT score) was retained for subsequent analyses^25^. For Nop effectors, structural units were delineated from the top-ranked models as contiguous regions of at least 15 amino acids with a pLDDT score over 70. Structural similarity between individual structural units was then evaluated with Foldseek^26^, using a normalised template modelling (TM)-score. Pairs of units with a TM-score ≥ 0.5 were considered structurally similar. The resulting similarity matrix was converted into a network and analysed with the R package igraph (https://igraph.org/r). Community detection across all structural units was performed using the Louvain algorithm^56^ (S4 Fig.). Structural visualisation and preparation of structure images were performed using UCSF ChimeraX^57^.

### Sequence- and structure-based functional annotation

Sequence-based functional annotation was performed with InterProScan^58^ against the Pfam database^59^. For each structural unit model, structural similarity searches were carried out with Foldseek^26^ against the Alphafold/Proteome, Alphafold/Uniprot50, Alphafold/Swiss-Prot and PDB100 databases. An additional structural similarity search was carried out for SU2 using DALI server^35^ against Alphafold/Arabidopsis proteome and Protein Data Bank (PDB) data bases^60,61^. The top five hits for each model were examined, and matches with a TM-score ≥ 0.5 were retained as putative structural and functional references. Finally, the similarity of SU6 and SU2 to RNA-dependent RNA polymerase (RdRP) module and to B3 and PUA domains, respectively, was further confirmed using the TM-align web server^62^.

### Subcellular localisation and prediction of transcriptional activation domains

Subcellular localisation was predicted using the plant cell–trained, machine learning–based tool LOCALIZER v1.0.4^42^, with effector sequences analysed as full-length proteins. Acidic transcriptional activation domains were predicted using two independent deep learning based methods, PADDLE (Predictor of Activation Domains using Deep Learning in Eukaryotes), trained on eukaryotic activation domains, and TADA (Transcriptional Activation Domain Activity), trained on plant specific activation domains^43,44^. For PADDLE, intrinsic structural disorder was first estimated for each effector with IUPred2A^63^, then used to compute activation scores and Z-scores for all overlapping 53 amino acid tiles spanning each protein. Regions comprising at least five consecutive tiles with Z-scores ≥ 4 or ≥ 6 were annotated as medium strength or high strength activation domains, respectively. TADA predictions were obtained using the TADA_T2 implementation provided as a Google Colab workflow (https://github.com/ryanemenecker/TADA_T2). For each effector, TADA scores were computed for all overlapping 40 amino acid tiles, and a per residue score between 0 and 1 was calculated as the mean of the scores for all tiles containing that residue (Table S8). In our study, a region was annotated as a putative activation domain only when the same region was predicted as activating by both PADDLE and TADA.

### Sequence analysis

Protein sequence alignments were generated with Clustal Omega^64^ and visualised using ESPript 3 (https://espript.ibcp.fr/ESPript). Evolutionary conservation profiles for SU1 and SU2 were generated with ConSurf-DB^65^ using the PDB_MSA pipeline on a Google Colab implementation (consurf.ipynb). For each representative SU1 or SU2 structure, residue conservation was estimated with a Bayesian method from structure guided multiple sequence alignments generated with FoldMason^66^, following a model selection pretest to identify the best fitting evolutionary substitution model.

### Identification and conservation analysis of putative activation domains in Nop homologues

Homologues of the 15 Nop effectors predicted to harbour both a putative nucleic acid–interaction module and a putative transcriptional activation domain were retrieved using phmmer searches implemented in HMMER v3.4^67^. Hits were retained using a minimum threshold of 40% amino-acid identity over at least 70% of the query length. These searches were conducted against 654 proteomes, including 417 from the genus Bradyrhizobium, 200 from Sinorhizobium/Ensifer, and 37 from Mesorhizobium, and yielded 643 non-redundant homologous proteins. Each sequence was analysed independently with PADDLE as described above, and homologues were then aligned using Clustal Omega. For every alignment position, the fraction of sequences with a PADDLE Z-score ≥ 4 was computed, thereby generating a position-wise conservation profile. Finally, a predicted activation domain was considered conserved when at least five consecutive alignment positions were predicted with a Z-score ≥ 4 in more than 75% of the homologous sequences.

### Detection of Nop-derived structural units in non-rhizobia type III effectors library

A dataset of 1,210 known T3SS effector sequences was compiled from multiple public databases, spanning 21 genera and including effectors from plant pathogenenic as well as animal-associated bacteria (S9 Table). Structural models for all effector sequences were predicted with AlphaFold as described above, and the top-ranked model for each effector was retained for downstream analyses. Each of the 22 SUs identified in Nop effectors was then used as an individual structural query to search this effector model set using Foldseek^26^. An SU was considered conserved when the corresponding Foldseek match reached a TM-score ≥ 0.75. For effectors with positive matches, SU conservation was further confirmed by manual structural superposition in UCSF ChimeraX.

### Usage of generative AI and AI-assisted technologies

We used ChatGPT-5.2 from OpenAI (https://chatgpt.com) during manuscript preparation to assist with English language editing, including correction of grammar, refinement of wording and occasional shortening of paragraphs. All outputs generated with this tool were carefully checked, revised when necessary, and approved by the authors, who retain full responsibility for the content of this publication.

## Acknowledgements

The authors would like to thank Philip Carella for compiling and sharing the plant-pathogenic bacterial effector sequences used in this study. This work was funded by the Gatsby Charitable Foundation (AT, SS, GAT3731/GLD).

## Competing interests

The authors declare that they have no competing interests.

## Author contributions

A.T. conceptualised the study. A.T. and S.S. designed, performed and interpreted analyses, wrote and edited the manuscript.

## Data and materials availability

All data needed to evaluate the conclusions in the paper are present in the paper and/or the Supplementary Materials. The best structural models for all bacterial type III effectors analysed in this study and the structural units extracted from Nop effectors have been deposited in Zenodo^25^ (https://doi.org/10.5281/zenodo.19162152).

## Supporting Information

**S1 Table. Nop effector dataset used in this study.**

List of Nodulation Outer Protein (Nop) effectors analysed in this study, indicating the bacterial strain of origin, protein identifiers from Microscope MaGe (mage.genoscope.cns.fr) and NCBI (ncbi.nlm.nih.gov) databases, effector family assignment, known function, and the references describing each effector.

**S2 Table. Sequence- and structure-based domain annotations of Nop effectors.**

Summary of the domain architecture of the Nop effectors analysed in this study. For each protein, the table indicates the positions in the amino-acid sequence of all domains identified by InterProScan and by structural unit segmentation (SU1 to SU22), together with their proposed functional annotation and the pLDDT score of the best AlphaFold model.

**S3 Table. Foldseek TM-scores for all pairwise structural comparisons between Nop-derived structural units identified in this study.**

All TM-scores resulting from the pairwise structural comparisons done by Foldseek of each Nop-derived structural unit compared to the others.

**S4 Table. Structural community composition analyses of the Nops-derived structural units.**

Communities identified by the Louvain community detection method are sorted by community size starting with the largest community. When possible annotations are provided.

**S5 Table. Foldseek matches of Nop-derived structural units.**

For each Nop structural unit, the five best structural hits detected by Foldseek are listed for the AlphaFold/Proteome, AlphaFold/UniProt50, AlphaFold/Swiss-Prot and PDB100 databases. For every match, the target protein, TM-score and Foldseek probability are reported.

**S6 Table. Dali-lite matches of the structural unit 2.**

The five best structural hits detected by Dali-lite are listed for the PDB and Alphafold/Arabidopsis proteome databases.

**S7 Table. Predicted nuclear localisation signals (NLS) in Nop effector and related effectors.**

LOCALIZER predictions of nuclear localisation signal (NLS) sequences for each effector analysed in this study. Up to five NLSs are predicted per effector protein sequence. for each. The start and end positions in the amino-acid effector sequence, as well as the predicted NLS motif, are reported.

**S8 Table. Transcriptional activation scores predicted by PADDLE and TADA.**

This table reports, for each amino-acid position in the effector sequences analysed, the PADDLE activation score, the corresponding PADDLE z-score, and the TADA activation score. Residues with PADDLE z-score ≥ 4 (medium-strength) and ≥ 6 (high-strength) are highlighted in blue and red, respectively, and residues with TADA score > 0.4 are highlighted in pink. A region is considered a putative activation domain when at least five consecutive residues are predicted as activating by both PADDLE and TADA.

**S9 Table. Catalogue of non-rhizobial type III effector structures used for comparative analyses.**

List of the 1,207 type III effector variants from plant-pathogenic and animal-associated bacteria. For each effector, the genus, species and effector name are given, together with the corresponding PDB file name as used in Dataset S3.

**S10 Table. Foldseek TM-scores for structural comparisons of Nop structural units with non-rhizobial type III effectors.**

Matrix of TM-scores computed with Foldseek for pairwise structural comparisons between the 96 structural domains extracted from Nop effectors and the 1,207 type III effector variants from plant-pathogenic and animal-associated bacteria.

**S11 Table. Summary of structural matches of Nop-derived structural units in non-rhizobial type III effectors.**

For each of the 22 Nop-derived structural units, the table reports the number of non-rhizobial effector variants from the set of 1,207 plant-pathogenic and animal-associated type III effectors with a significant structural match with TM-score ≥ 0.75.

## Supplementary Materials

## Supplementary Figures

**S1 Fig.**
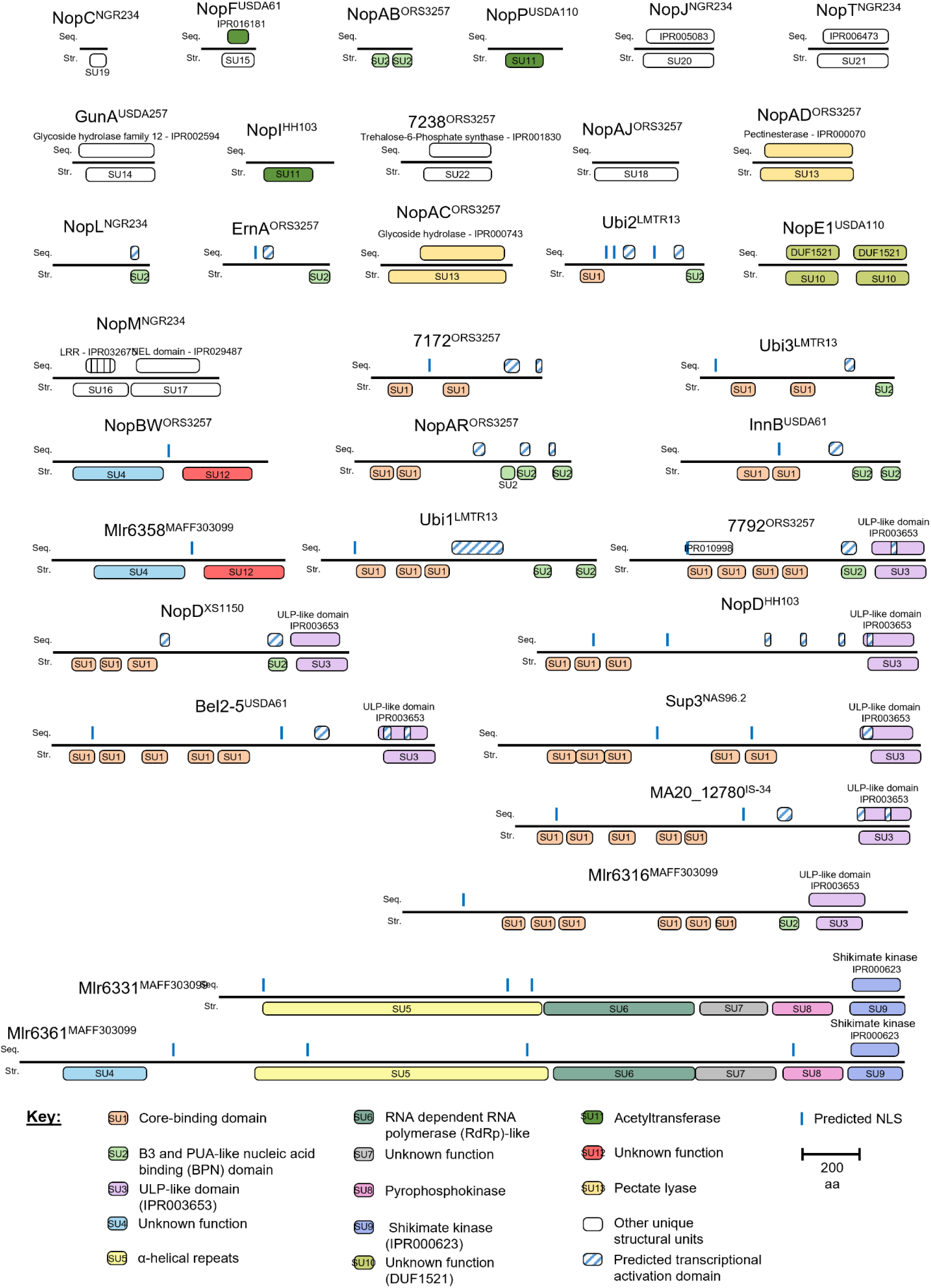
Modular domain architecture of rhizobial type III Nop effectors. Linear domain organisation of the 33 Nodulation Outer Protein (Nop) effectors analysed in this study. Each protein is shown as a horizontal bar, with sequence-derived domains displayed above the bar (seq.) and structural units (SU1 to SU22) defined by AlphaFold-based clustering displayed below (str.). For sequence-derived domains, the corresponding InterProScan accession numbers (IPR identifiers) are indicated above each domain. The identity of the different displayed domains and sequence motifs is indicated in the key. Only regions identified as activation domains by both PADDLE and TADA are shown as transcriptional activation domains.

**S2 Fig.**
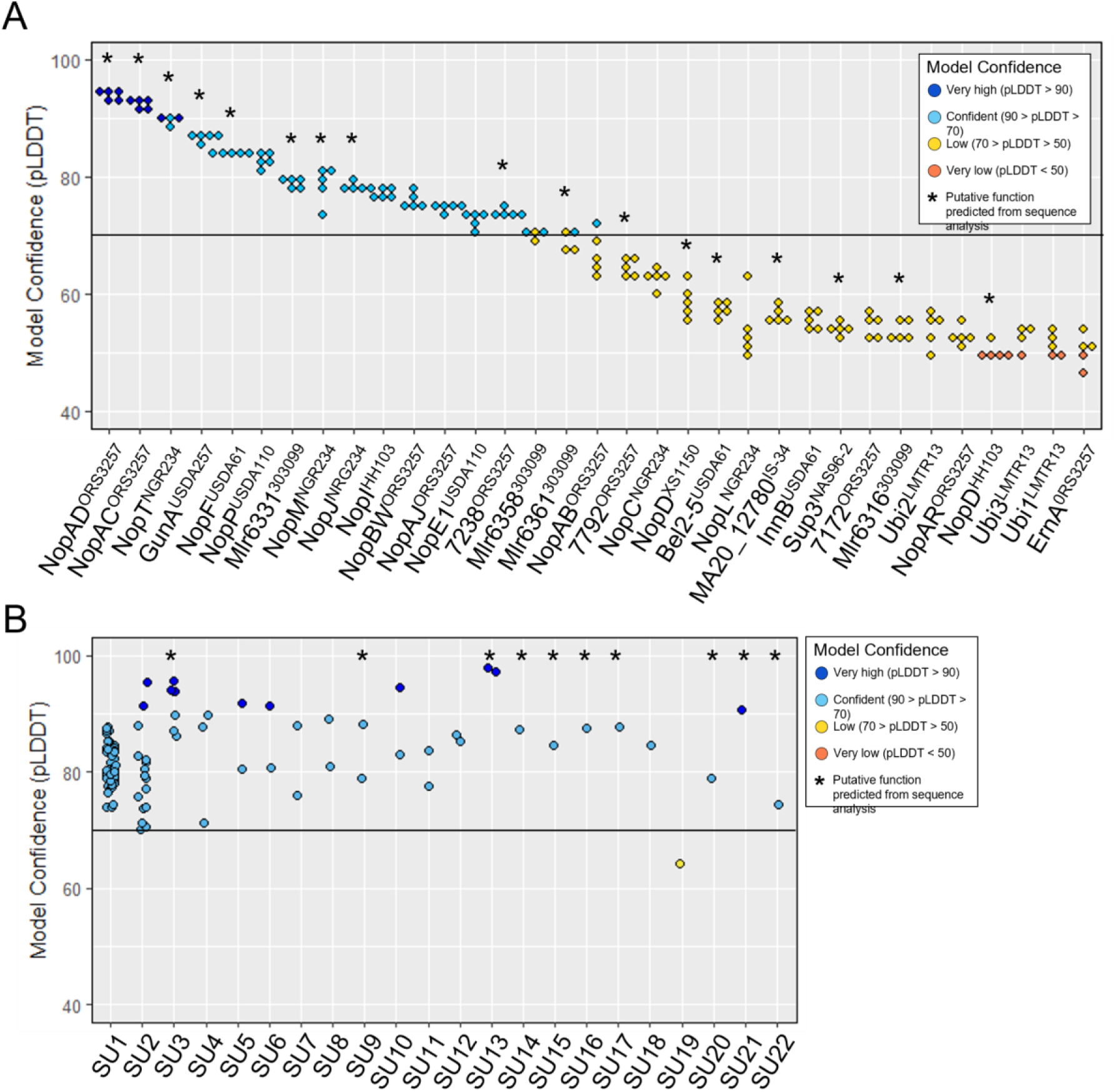
AlphaFold confidence scores for full-length Nop effector models and derived structural units. (A) For each effector, five full-length models were generated by AlphaFold and ranked from highest to lowest confidence based on the overall predicted local distance difference test (pLDDT) score. The top-ranked model was retained for subsequent structural analyses. (B) pLDDT scores for the individual structural units segmented from the full-length Nop models. For both panels, the horizontal line marks the pLDDT = 70 threshold separating low- from confident-quality predictions, and points are coloured by the standard AlphaFold confidence classes. Asterisks indicate effectors for which at least one putative functional domain was detected by InterProScan.

**S3 Fig.**
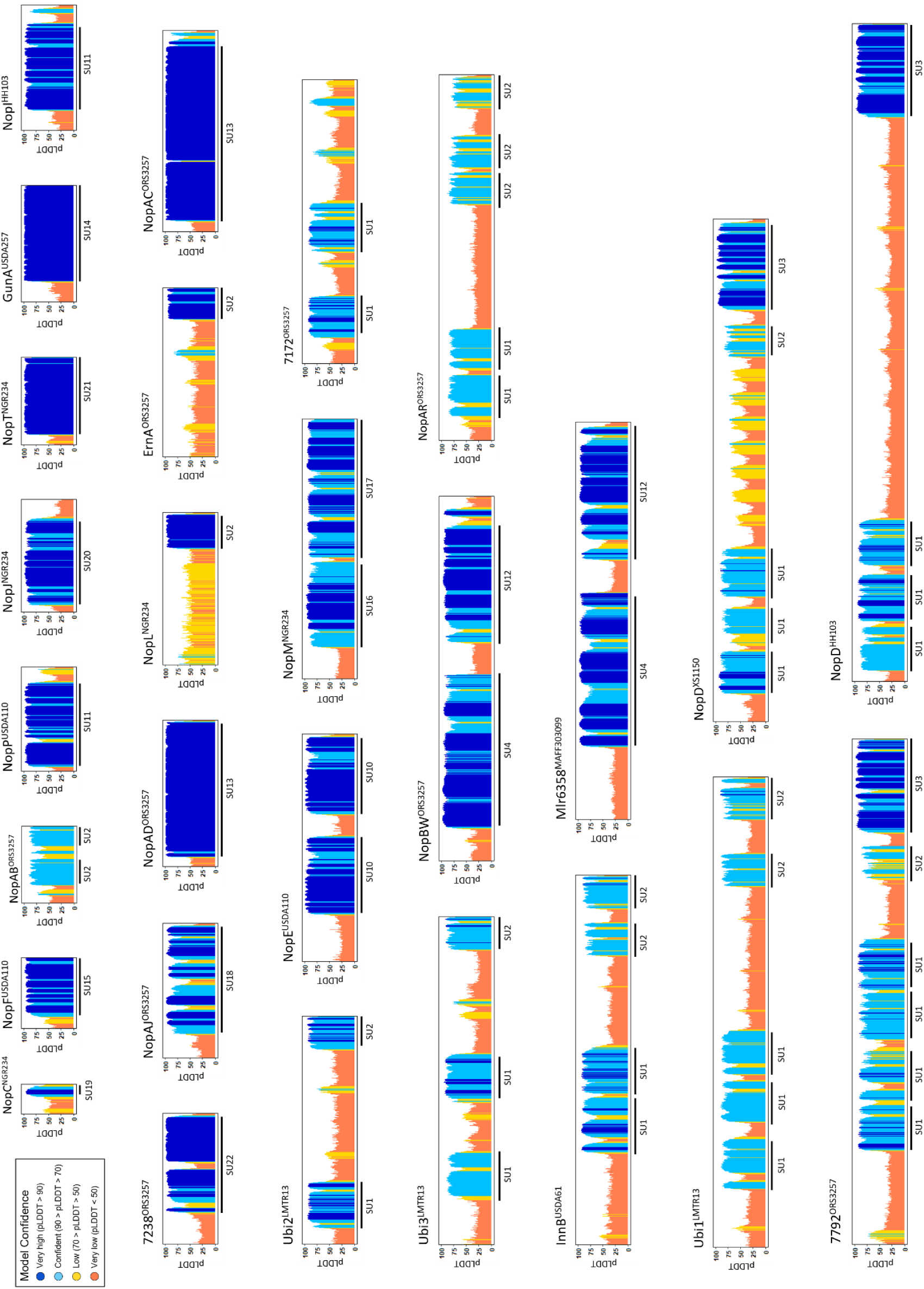
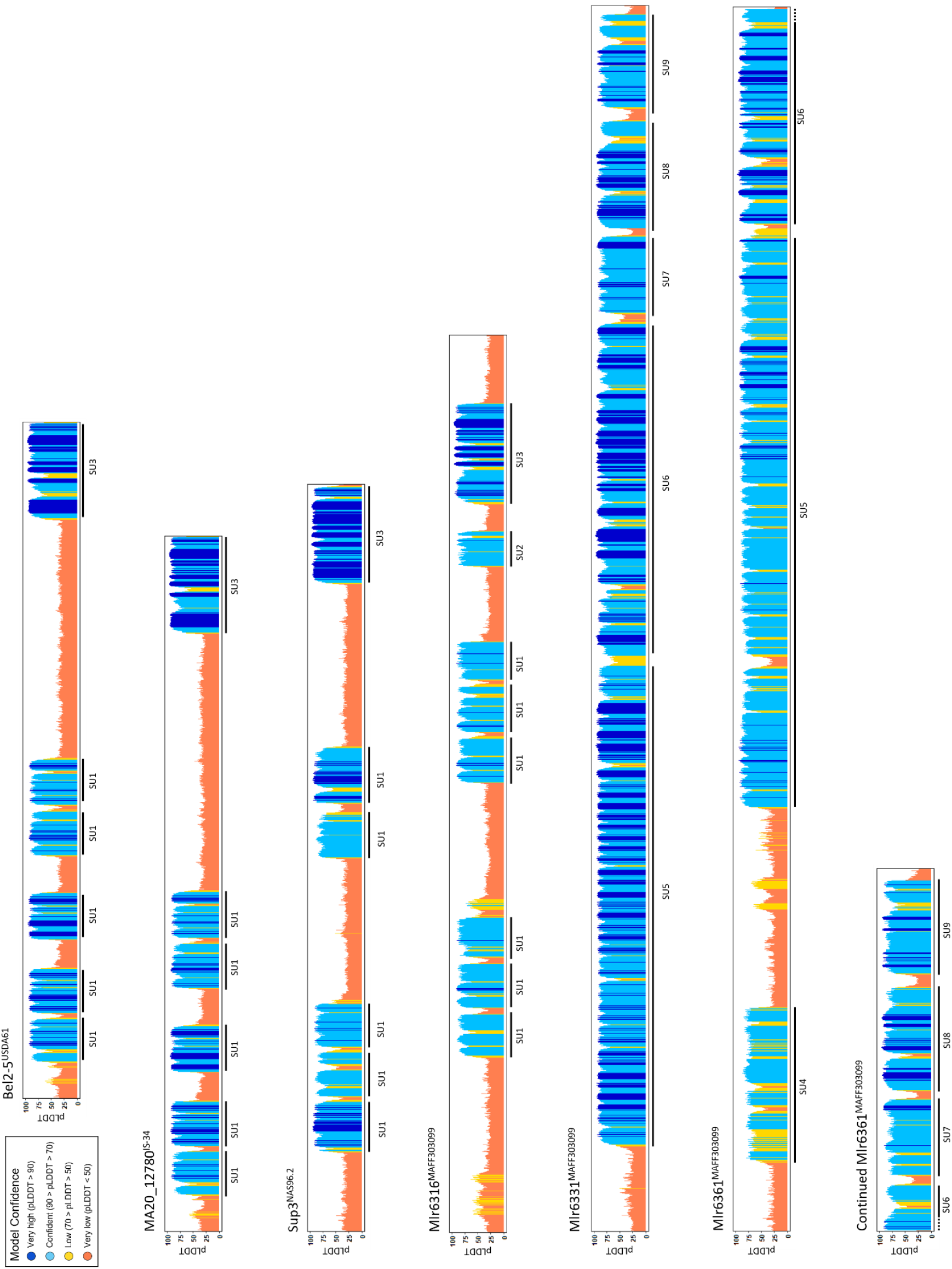
Structural unit identification using local model prediction confidence. Diagrams of per-residue predicted local distance difference test (pLDDT) values for all Nop effectors analysed in this study. Structural units (SU1 to SU22) corresponding to contiguous regions with high pLDDT values (≥70) are underlined. The identity of each structural unit, as defined by structure-based clustering, is indicated below the line.

**S4 Fig.**
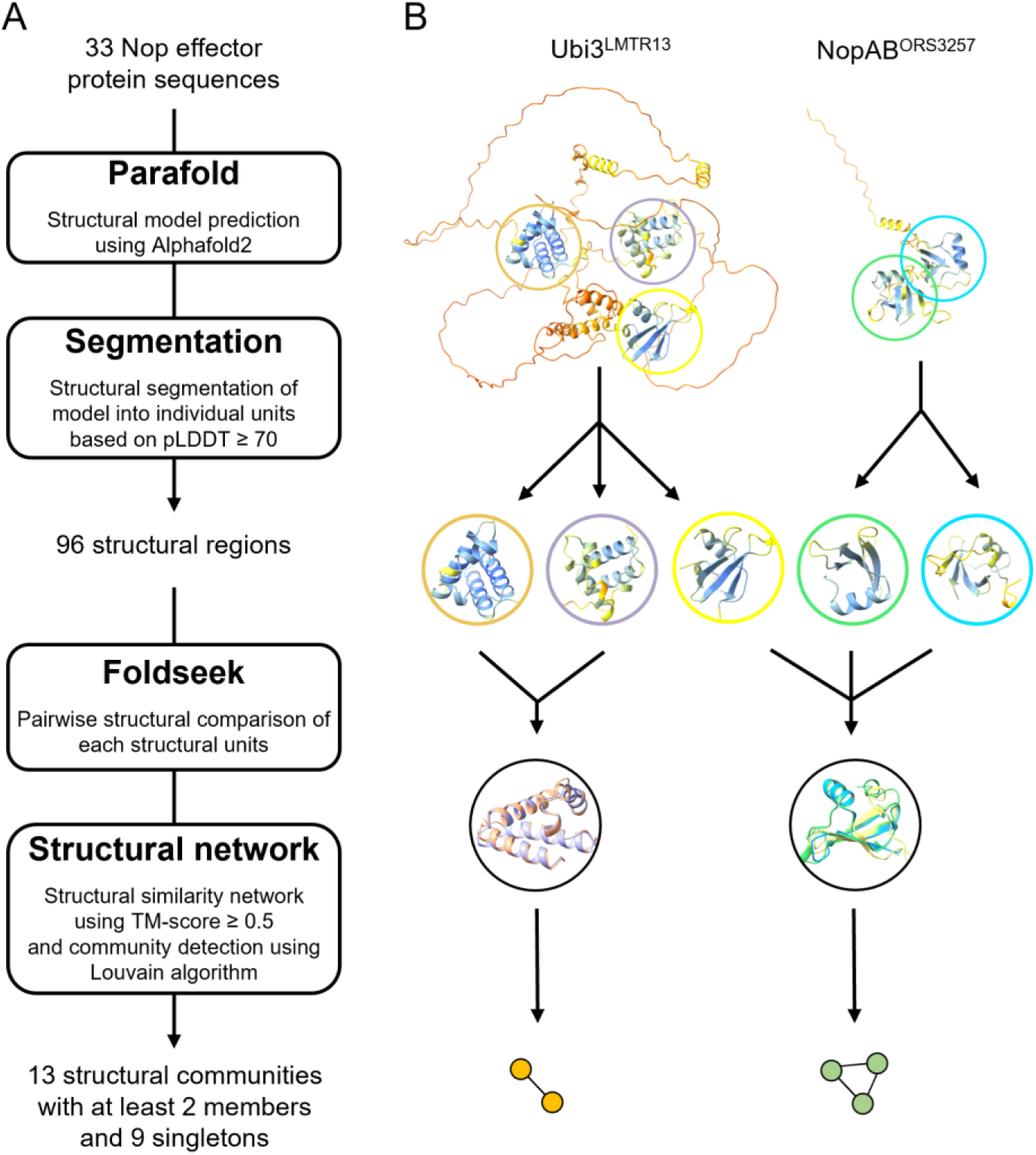
Workflow for structure-based analysis of Nop effectors. (A) Overview of the computational pipeline used to analyse Nop effector structures. (B) Example illustrating the segmentation and structural clustering of two Nop effectors.

**S5 Fig.**
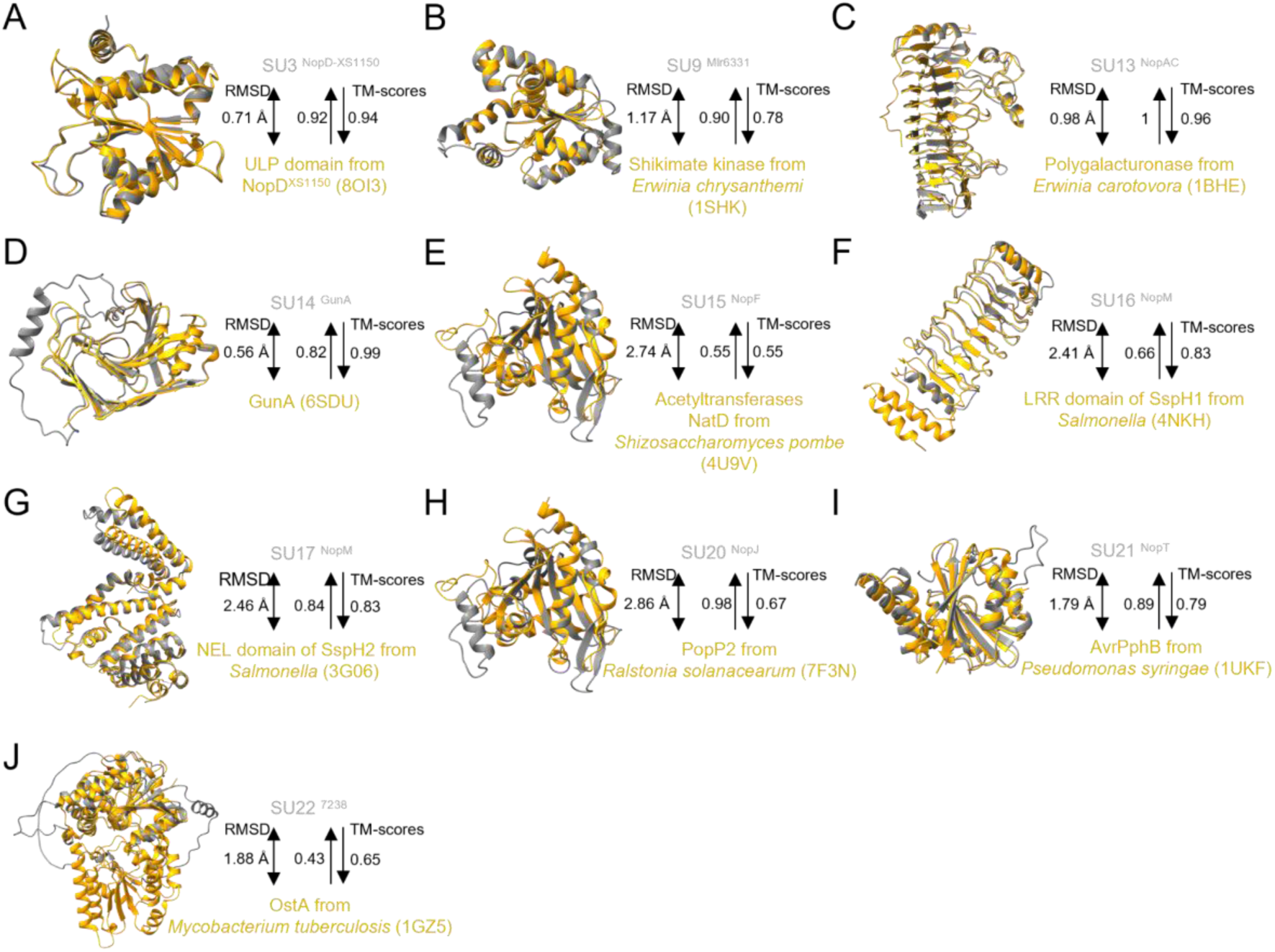
Structural similarity between annotated Nop structural units and crystallographic reference folds. Structural superpositions between Nop-derived structural units (SUs) for which a sequence-based annotation was identified (grey) and representative crystallographic reference structures (yellow). Reference PDB entries used for each comparison are indicated in brackets. Structural similarity between SU models and reference structures is supported by low root-mean-square deviation (RMSD < 3 Å) and high template modelling scores (TM-score ≥ 0.5), indicating that the structural segmentation of Nop recaptures the expected crystallographic folds.

**S6 Fig.**
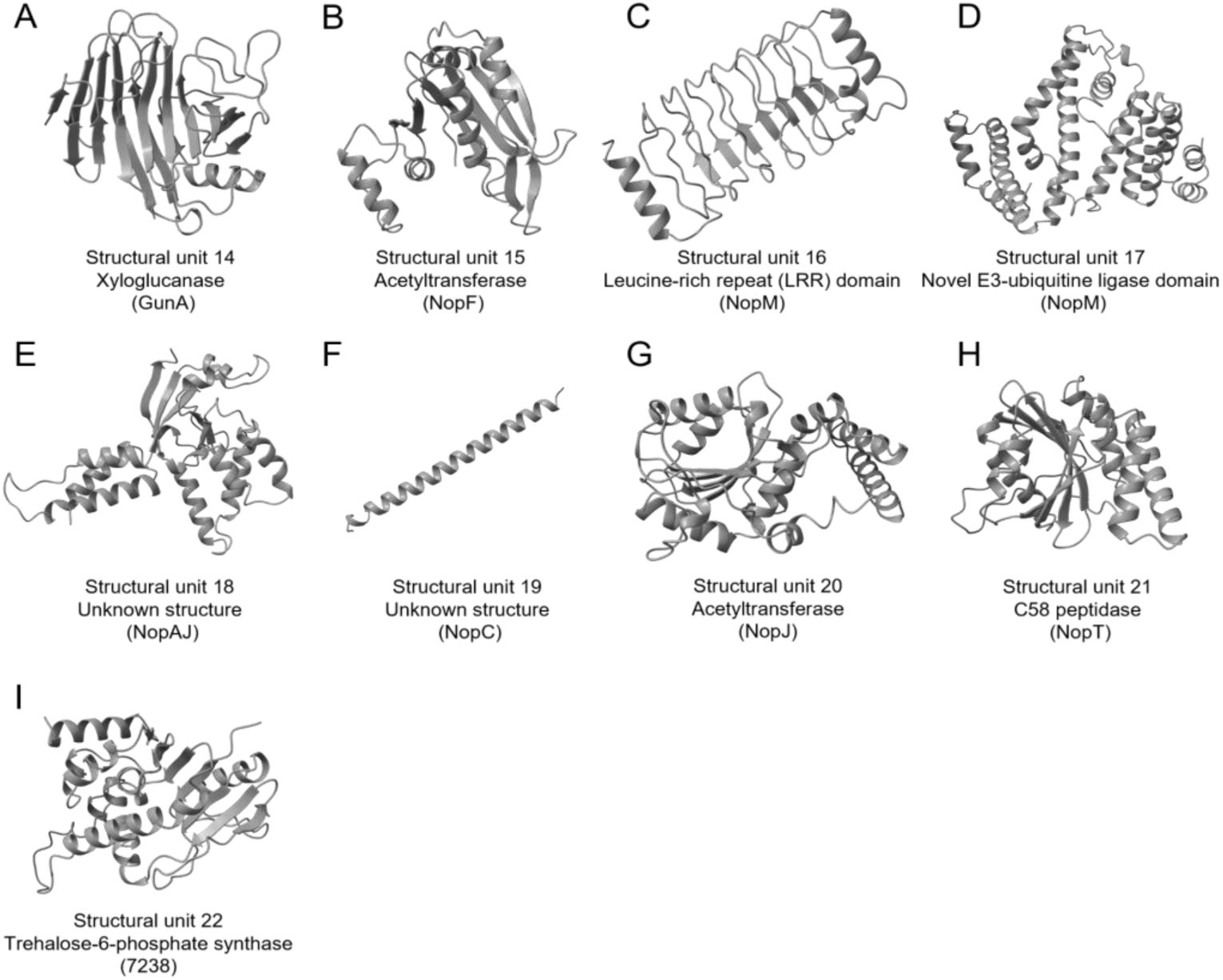
Unique structural units identified as singletons in the Nop effector repertoire. Each panel shows an AlphaFold model of a structural unit that formed a singleton node in the structural similarity network. These units did not cluster with any other structural region at a TM score above 0.5. The putative fold or function inferred from Foldseek analysis is indicated below each panel, with the corresponding Nop effector in which the unit is found indicated in parentheses.

**S7 Fig.**
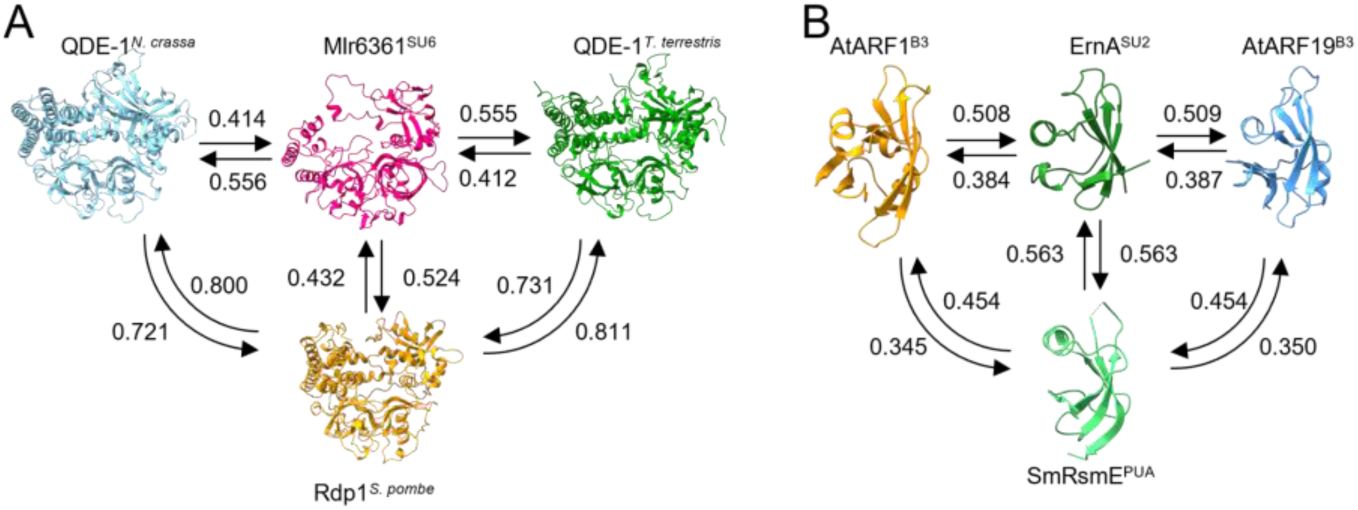
Structural alignment of SU6 and SU2 with their closest matches. Structural comparisons of Nop structural units SU6 and SU2 with their closest structural neighbours using TM-align. TM-scores calculated in each direction of the alignment are shown, and arrows indicate the direction of the corresponding TM-align comparison. (A) Superposition of SU6 from Mlr6361MAFF303099 with RNA-dependent RNA polymerase domains from Qde-1 of Thermothielavioides terrestris (PDB: 5FSW) and Neurospora crassa (PDB: 7Y7R), and Rdp1 from Schizosaccharomyces pombe (Alphafold model: AF-O14227-F1). (B) Superposition of ErnA SU2 with the B3 domains of ARF1 (PDB: 4LDX) and ARF19 (Alphafold model AF-Q8RYC8-F1) from Arabidopsis thaliana and the PUA RNA-binding domain of RsmE (PDB: 4J3C) from Sinorhizobium meliloti. The BPN (SU2) domain model aligned most closely with the PUA domain.

**S8 Fig.**
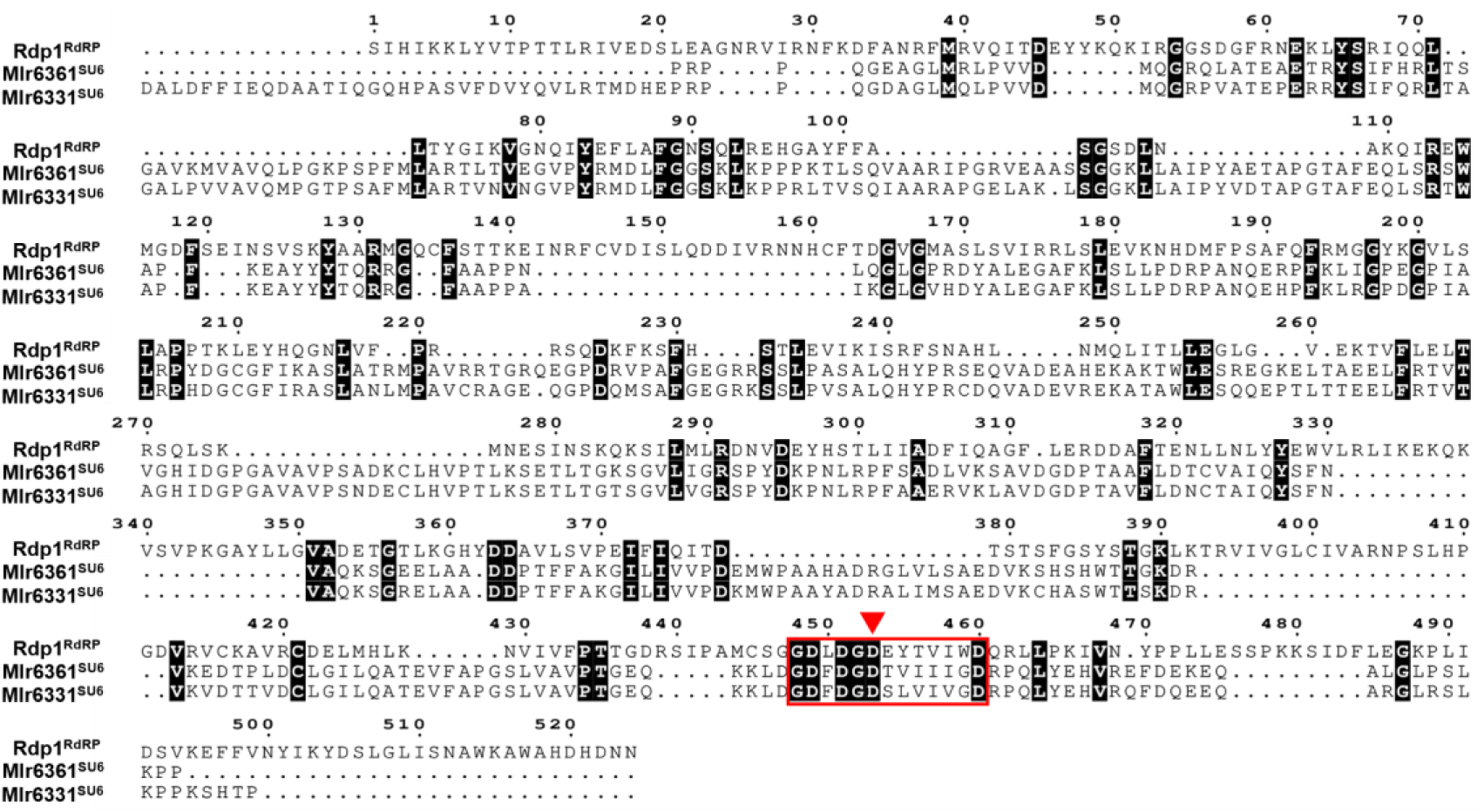
Limited sequence similarity between the structural units 6 and the two-barrel polymerase module of Rdp1. Multiple sequence alignment of the two-barrel RNA-dependent RNA polymerase (RdRP) module of Rdp1 from Schizosaccharomyces pombe (Uniprot accession: O14227) with the SU6 regions of Mlr6331MAFF303099 and Mlr6361MAFF303099. Overall sequence similarity is very low, with only short stretches of conservation, notably in the catalytic site of the polymerase (boxed in red), where the key aspartate residue (red triangle) known to be critical for the RdRP catalytic activity is conserved in the three sequences. Residues conserved in all 3 sequences are highlighted with a black background.

**S9 Fig.**
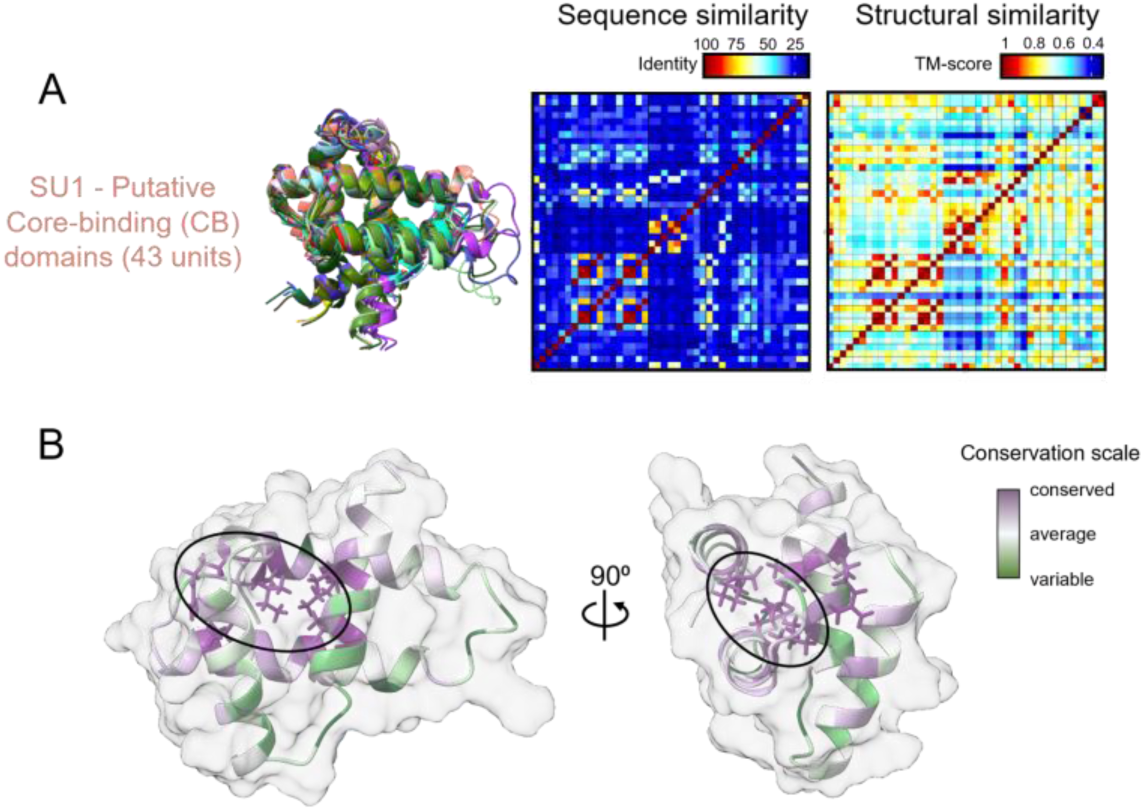
Sequence and structural features of the structural unit 1. Analysis of 43 occurrences of SU1 identified in Nop effectors. (A) The structural overlay of all SU1 units (left), the sequence-based similarity matrix (centre), and the structure-based similarity matrix derived from pairwise TM-scores (right), highlighting stronger conservation at the structural than at the sequence level. (B) ConSurf-based mapping of evolutionary conserved residues onto the SU1 structural model, highlighting the formation of an internal hydrophobic core (black circles).

**S10 Fig.**
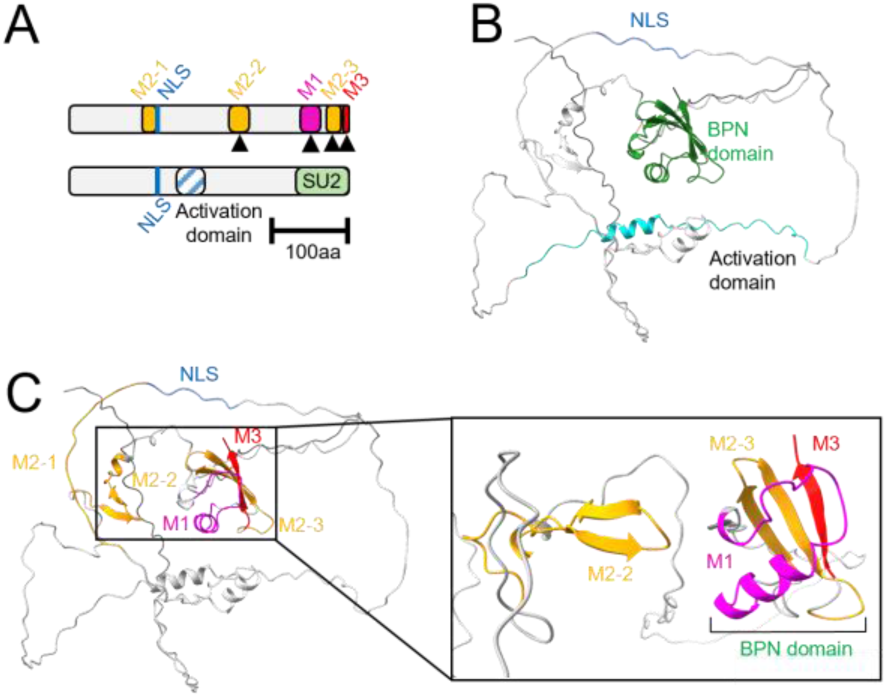
The BPN domain of ErnA aligns to experimentally validated activity-associated motifs. (A) Previously defined ErnA sequence motifs are shown above (M1, three M2 motifs, and M3) and aligned with the domain architecture inferred in this study shown below. Black triangles mark motifs experimentally demonstrated to be required for ErnA biological activity. (B) AlphaFold model of ErnA annotated with the domains defined in this study. (C) The M1 motif folds into an α-helix within the BPN domain, positioned opposite the β-sheet surface formed by motifs M2–3 and M3.

**S11 Fig.**
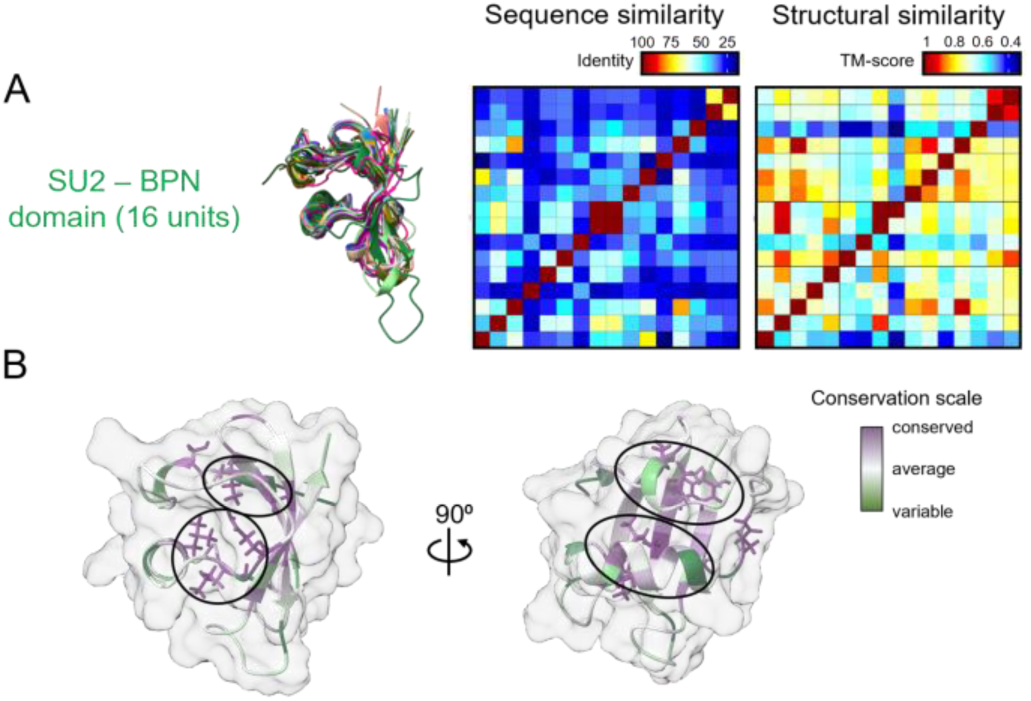
Sequence and structural features of the structural unit 2. The 16 occurrences of SU2 (B3 and PUA-like nucleic acid binding domain, BPN) from Nop effectors were compared at the structural and sequence levels. (A) The structural overlay of all SU2 units (left), sequence-based similarity matrix (centre), and structure-based similarity matrix derived from pairwise TM-scores (right), reveal stronger conservation of the overall fold than of the primary sequence. (B) Projection of ConSurf-based evolutionary conservation onto SU2 indicates that the highest conserved residues form an internal hydrophobic core (black circles).

**S12 Fig.**
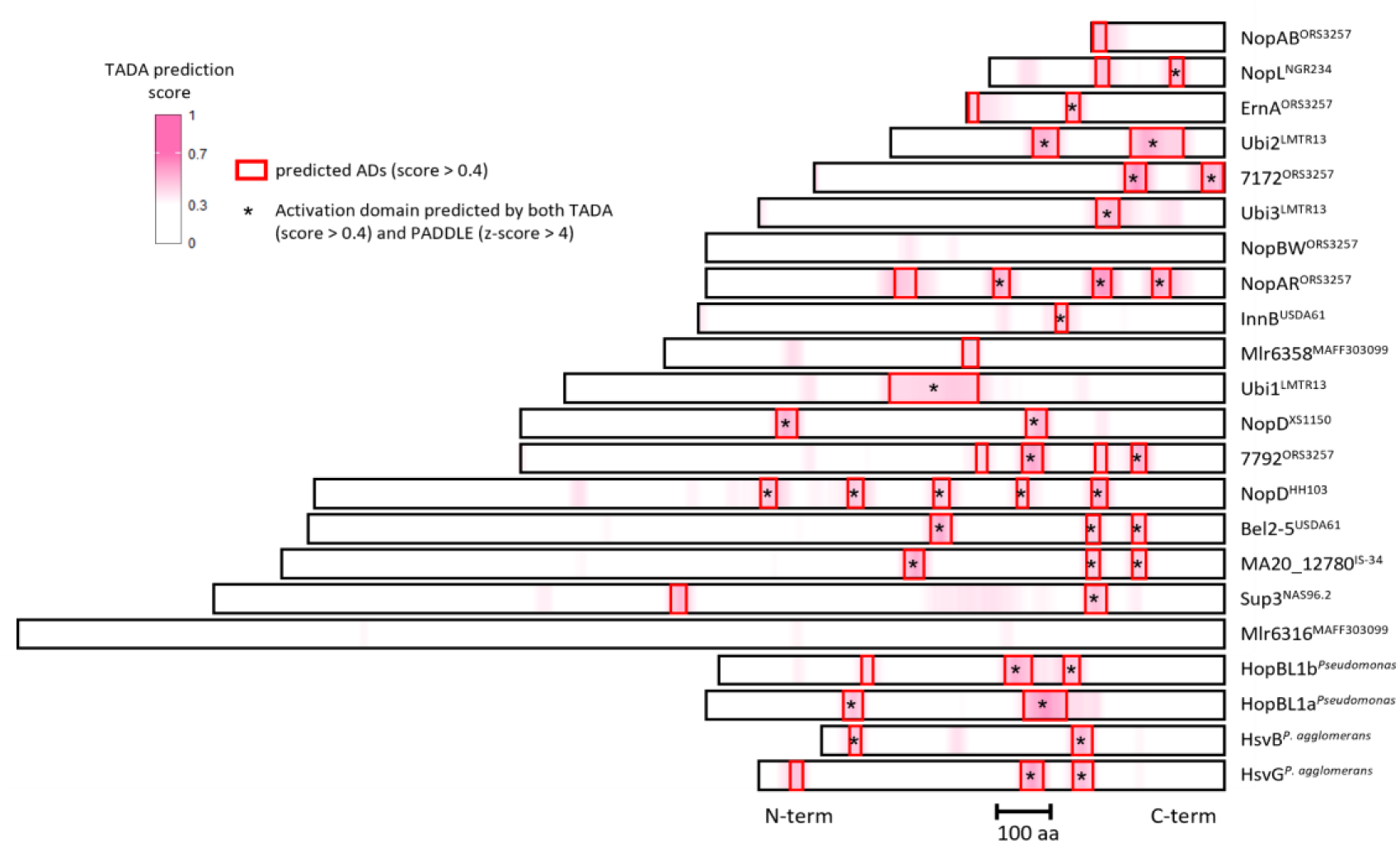
TADA-based prediction of transcriptional activation domains in Nop effectors. Heatmap of TADA prediction scores along Nop effector sequences, plotted from N-terminus to C-terminus. The colour scale indicates the local TADA activation score, with more intense shades corresponding to higher predicted activation potential. Red boxes highlight regions with a TADA score ≥ 0.4, the minimum threshold used to define predicted transcriptional activation domains. Black asterisks indicate activation domains predicted independently by both TADA and PADDLE.

**S13 Fig.**
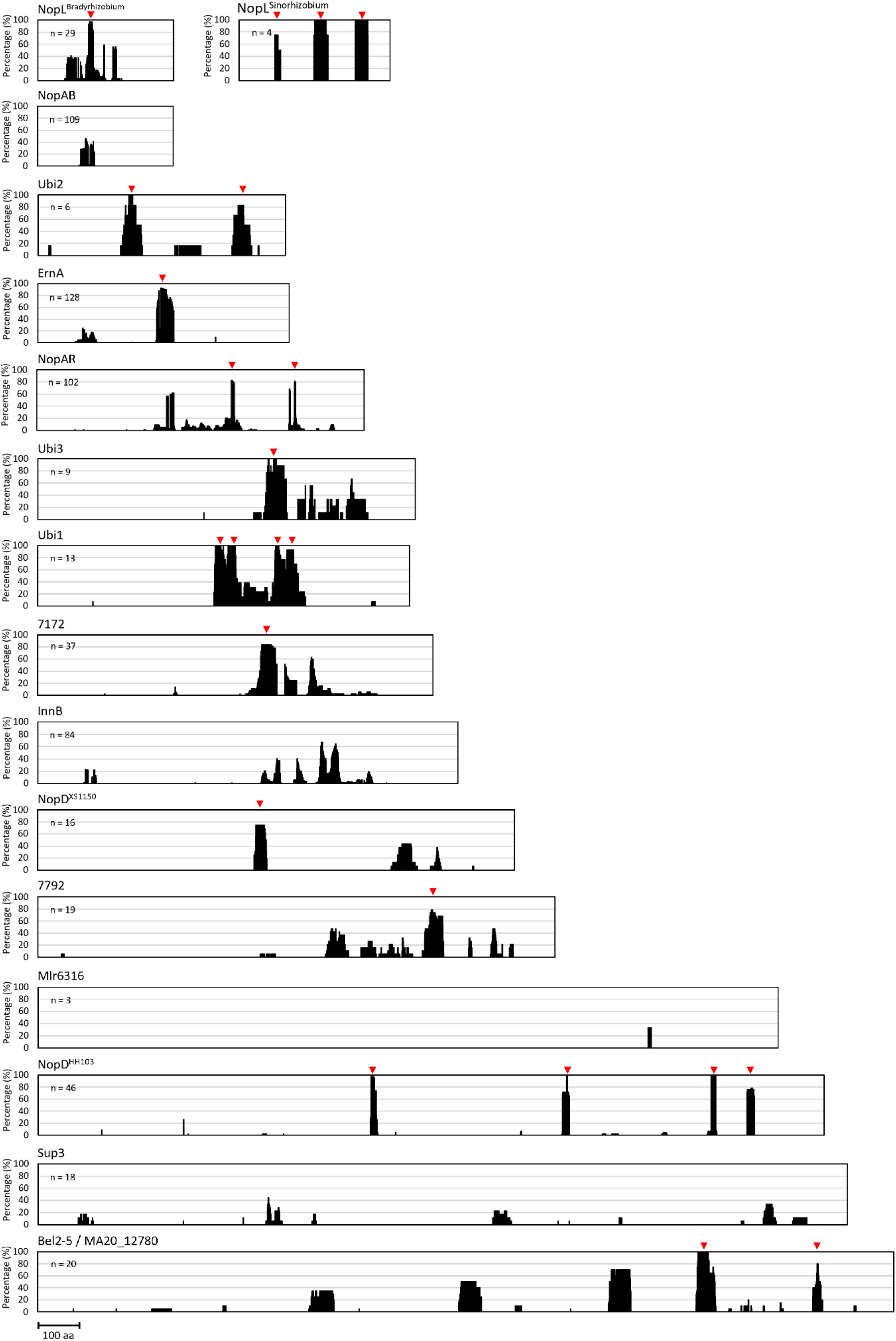
Conservation of predicted transcriptional activation domains in Nop effector homologues. For each Nop effector family carrying a predicted activation domain in this study, the plots show the proportion of homologous sequences with a PADDLE Z-score ≥ 4 at each position of the protein alignment. Bars indicate sites predicted as transcriptional activation domains across the alignment. Regions are considered as conserved transcription activation domains when more than 75% of homologues exceed the threshold (red triangles). n indicates the number of homologous proteins analysed for each effector family.

**S14 Fig.**
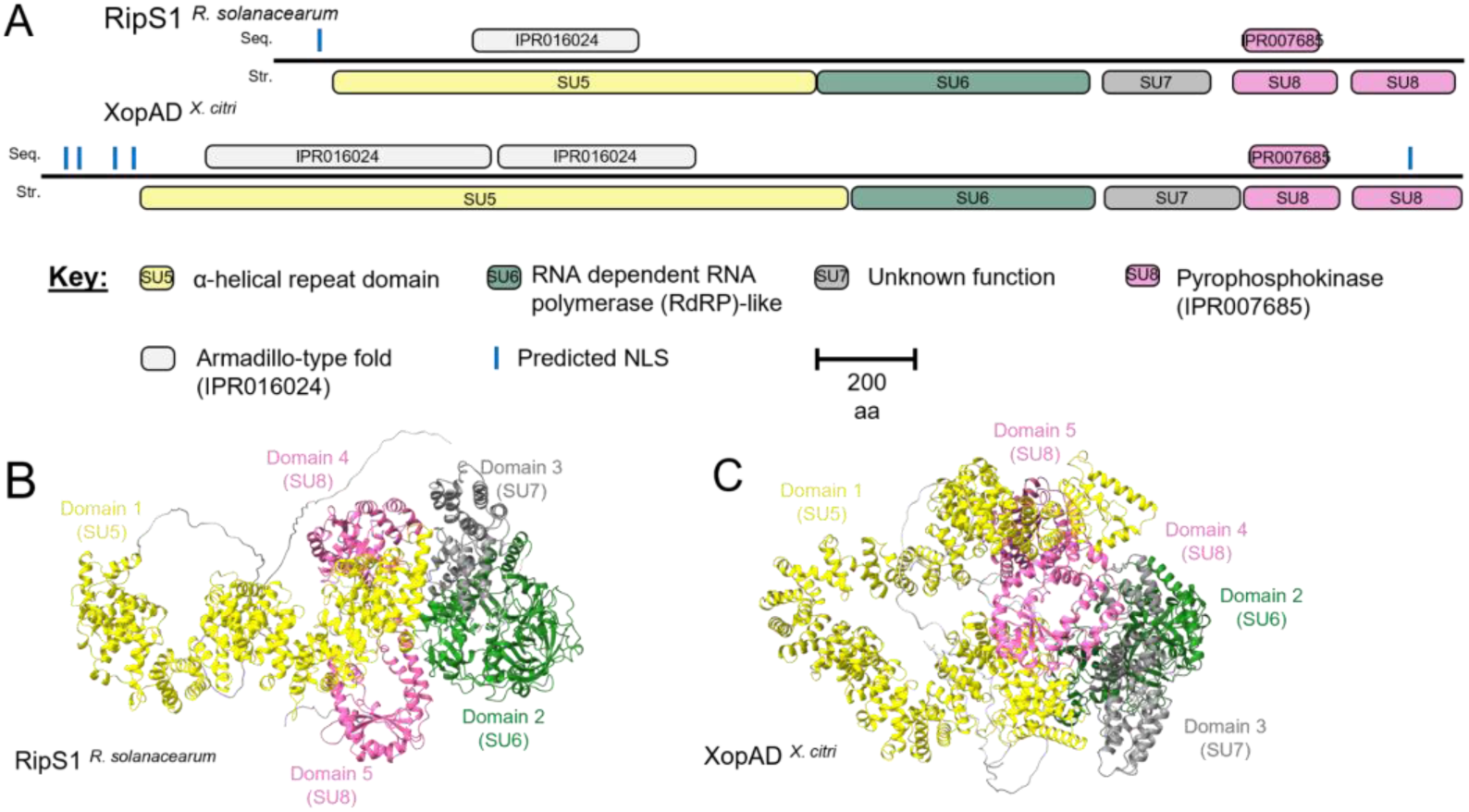
Modular architecture of SKPW-family effectors RipS1 and XopAD. (A) Linear domain organisation of the SKPW-family effectors RipS1 from Ralstonia solanacearum and XopAD from Xanthomonas citri. Each protein is shown as a horizontal bar, with sequence-based annotations (InterProScan domains; IPR identifiers indicated above) displayed above the bar (seq.) and structural units defined in this study displayed below (str.). (B,C) AlphaFold models of RipS1 and XopAD, respectively, segmented into five structural units.

**S15 Fig.**
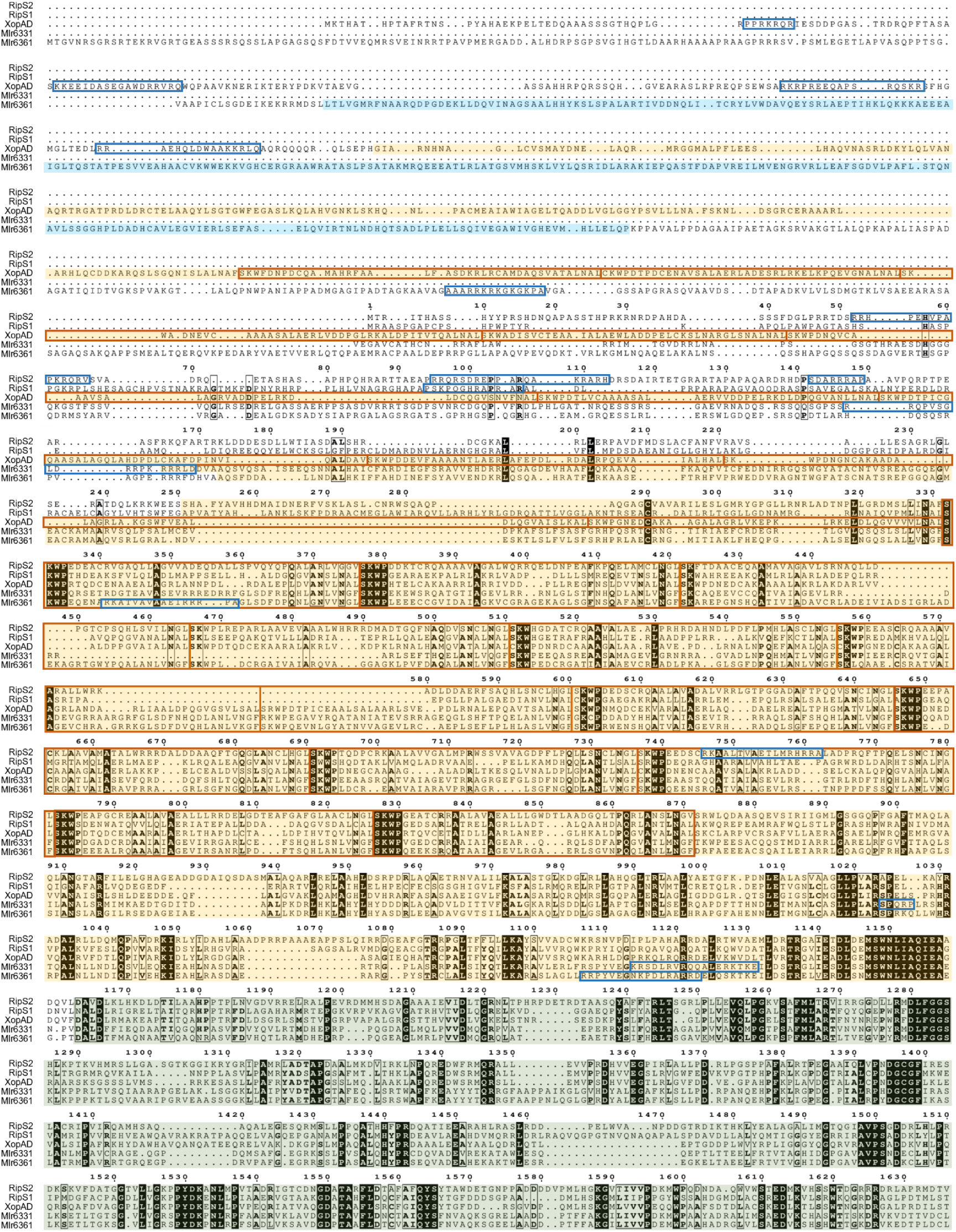
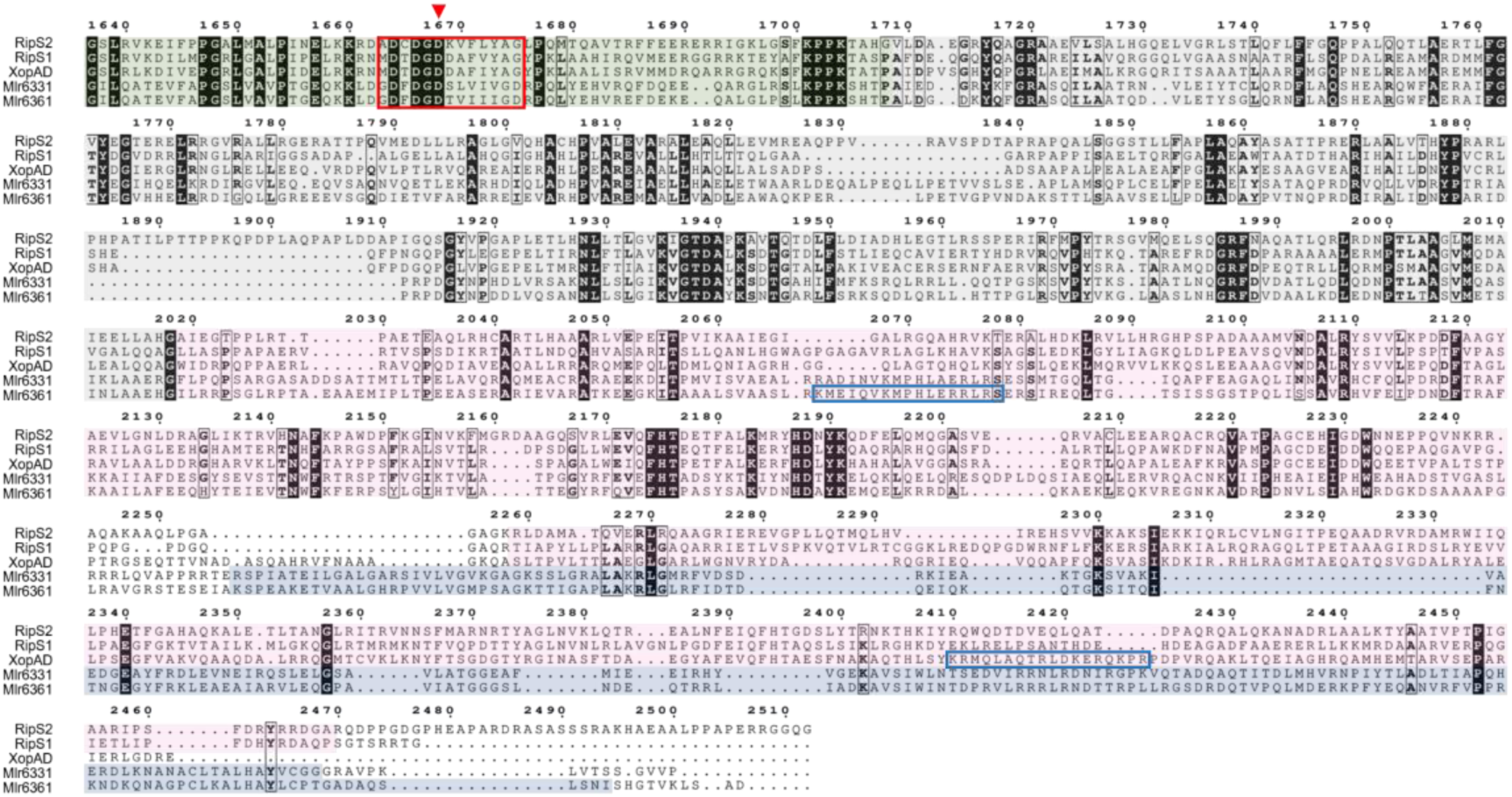
Multiple protein sequence alignment of SKPW-family effectors RipS1, RipS2, XopAD and the two Nop effectors Mlr6331 and Mlr6361. Although overall sequence identity is low, the regions corresponding to the predicted structural units are clearly conserved and highlighted along the alignment. The α-helical repeat unit SU5 (yellow), the RdRP-like unit SU6 (green), SU7 (grey) and the GTP pyrophosphokinase unit SU8 (pink) are present in all five effectors in the same order. A fifth unit, SU4 (clear blue), occurs only in Mlr6361, whereas the C-terminal shikimate kinase domain SU9 (dark blue), conserved in both Mlr6331 and Mlr6361, is replaced by a second SU8 unit in RipS1, RipS2 and XopAD. LOCALIZER predicts between one to five nuclear localisation signals (NLSs) in each effector (boxed in light blue). All sequences retain SKPW repeats (orange boxes), which overlap the SU5 α-helical repeat region. Within each RdRP-like SU6 module, residues around the predicted catalytic site are conserved (red box), including the key catalytic aspartate known to be essential for RdRP activity (red triangle). Residues conserved in 4 of the 5 effector protein sequences are outlined in black, whereas positions invariant in all five sequences are highlighted with a black background.

## Notes

https://doi.org/10.5281/zenodo.19162152

### Competing Interest Statement

The authors have declared no competing interest.

